# Temporin B forms hetero-oligomers with Temporin L, modifies its membrane activity and increases the cooperativity of its antibacterial pharmacodynamic profile

**DOI:** 10.1101/2022.03.09.483583

**Authors:** Philip M. Ferguson, Maria Clarke, Giorgia Manzo, Charlotte K. Hind, Melanie Clifford, J. Mark Sutton, Christian D. Lorenz, David A. Phoenix, A. James Mason

**Author notes:** These authors contributed equally (Philip M. Ferguson, Maria Clarke, Giorgia Manzo). **Corresponding Author**, A. James Mason, King’s College London, Institute of Pharmaceutical Science, Franklin-Wilkins Building 150 Stamford Street, London, SE1 9NH, UK.

## Abstract

The pharmacodynamic profile of antimicrobial peptides (AMPs) and their *in vivo* synergy are two factors that are thought to restrict resistance evolution and ensure their conservation. The frog *Rana temporaria* secretes a family of closely related AMPs, temporins A-L, as an effective chemical dermal defence. The antibacterial potency of temporin L has been shown to increase synergistically in combination with both temporins B and A but this is modest. Here we show that the less potent temporin B enhances the cooperativity of the *in vitro* antibacterial activity of the more potent temporin L against EMRSA-15 and that this may be associated with an altered interaction with the bacterial plasma membrane, a feature critical for the antibacterial activity of most AMPs. Addition of buforin II, a histone H2A fragment, can further increase the cooperativity. Molecular dynamics simulations indicate temporins B and L readily form hetero-oligomers in models of Gram-positive bacterial plasma membranes. Patch-clamp studies show transmembrane ion conductance is triggered with lower amounts of both peptides and more quickly, when used in combination, but conductance is of a lower amplitude and pores are smaller. Temporin B may therefore act by forming temporin L/B hetero-oligomers that are more effective than temporin L homo-oligomers at bacterial killing and/or by reducing the probability of the latter forming until a threshold concentration is reached. Exploration of the mechanism of synergy between AMPs isolated from the same organism may therefore yield antibiotic combinations with advantageous pharmacodynamic properties.

## Introduction

Host defence peptides (HDPs) are multifunctional molecules that are key components of the innate immune system and are found in all classes of life. There is interest in developing HDPs for therapeutic use as part of the response to the global increase in antimicrobial resistance, with some peptides able to combat infections by influencing the host immune response and antimicrobial peptides (AMPs) possessing highly potent bactericidal activity.^1^ Because of this and because, unlike clinically relevant antibiotics, HDPs and AMPs are produced by metazoans to counter infections, there is also interest in understanding how they have remained effective throughout evolutionary history.^2^

In laboratory conditions, serial passage of e.g. *Staphylococcus aureus* in the presence of AMPs leads to reduced susceptibility to both clinically prescribed antibiotics and human HDPs and this can be achieved with no detectable impact on fitness.^3^ An alternative perspective however is provided by work that has shown that, while adaptation to AMPs is indeed readily achievable, the resulting resistance levels are generally far lower than obtained with antibiotics under the same conditions.^4^ The modifications that arise as a result of bacterial adaptation to AMPs include changes in membrane surface charge, potential, permeability and fluidity, and the production of outer membrane vesicles and these may lead to altered susceptibility to HDPs, a reduction in host colonization and increased stimulation of host macrophages.^2^ It remains possible therefore that, as has been shown for *mcr-1*,^5^ the trade-offs required between fitness and resistance are such that the resistance that is achievable against AMPs has a ceiling below that observed for antibiotics, in particular in an *in vivo* setting.

It is further suggested that the pharmacodynamics of AMPs reduces the probability of resistance emerging.^6^ Antimicrobial agents that have a more cooperative, dose dependent activity, as characterised by a steeper slope in a dose-response curve, benefit from a narrow mutant selection window which results from a smaller concentration range where efficacy is incomplete. AMPs have, in general, a more cooperative dose-response than antibiotics and hence a smaller window in which a selective pressure will be exerted.^7^ The chemical composition of the infection setting may play an important role in limiting the ability of pathogenic bacteria to adapt to the innate immune response by ensuring that multiple AMPs, with differing mechanisms of action, are available. Cross-resistance between AMPs with differing mechanisms of action has been shown to be low,^8^ while combining AMPs with differing mechanisms of action, from different organisms, has been shown to further enhance the cooperativity of the dose-response^7^ and hence the pharmacodynamic properties of combinations of AMPs may further limit the risk of resistance emerging. The extent to which combinations of AMPs from the same organism act in synergy and how they might interact to produce a more cooperative dose-response is however yet to be fully explored.

The temporins comprise a very well-studied family of AMPs that now number more than 130 peptides,^9^ and whose members have been extensively evaluated and engineered to gain superior antibacterial activity.^10^ Temporin L is a broad-spectrum AMP, with potent bactericidal activity, identified, along with nine further temporins, in the European red frog *Rana temporaria*.^11^ Synergy against Gram-negative species has been described between temporin L and each of temporin A and temporin B which, individually, have only weak activity.^12^ Temporin L was shown to disrupt homo-oligomerisation of both temporin A and temporin B, behaviour that would enhance their translocation across the outer membrane and access the bacterial plasma membrane, the presumed site of their membrane disruptive activity.^12^ Temporins A and L differ substantially in their molecular mechanisms of action,^13^ and our previous work has identified fundamental differences in how temporins B and L insert into, and induce ion conductance in, models of the bacterial plasma membrane.^14,15^ Here we use two time-resolved biophysical methods - molecular dynamics (MD) simulations and patch-clamp - to examine how a combination of temporin B and temporin L inserts into and disrupts models of the Gram-positive plasma membrane. We use this understanding to explain how the less potent temporin B can influence the cooperativity of the dose dependent bactericidal activity of temporin L against methicillin resistant *S. aureus*, which is in-turn compared with that of existing, clinically relevant antibiotics. Together, this provides a mechanistic perspective of how AMPs from the same organism may combine to enhance the pharmacodynamic profile and consequently reduce the risk of resistance, to the innate immune response, emerging.

## Experimental procedures

### Peptides and lipids

Temporin L, temporin B, buforin II and pleurocidin were purchased from Cambridge Research Biochemicals (Cleveland, UK) as desalted grade (crude) and were further purified using water/acetonitrile gradients using a Waters SymmetryPrep C8, 7 μm, 19 × 300 mm column. All peptides were amidated at the C-terminus. The lipid 1,2-diphytanoyl-*sn*-glycero-3-phospho-(1’-*rac*-glycerol) (DPhPG) was purchased from Avanti Polar Lipids, Inc. (Alabaster, AL) and used without any purification. All other reagents were used as analytical grade or better.

### Antibacterial activity assay

The antibacterial activity of the peptides was assessed through a modified two-fold broth microdilution assay with modal MICs generated from at least three biological replicate experiments.^14–16^ The method broadly followed EUCAST methodology, with non-cation adjusted Mueller Hinton replacing cation-adjusted Mueller Hinton. Peptides and antibiotics were diluted in a two-fold dilution in media down a sterile, polypropylene 96 well plate (Greiner Bio-One GmbH, Frickenhausen, Germany). Bacteria were then added, back-diluted from an overnight culture, at a starting concentration of 5 × 10^5^ CFU/ml. Plates were incubated, static at 37 C, for 20 hours and the OD_600_ was determined using a Clariostar plate reader (BMG Labtech). The MIC was defined as the lowest concentration where growth was <0.1 above the background absorbance. For temporin B/temporin L synergy screening experiments, MICs were performed as above, but with molar ratios of the two AMPs, i.e. 1:1, 3:1 and 1:3 for temporin L:temporin B. To test for synergy between temporin L, temporin B and their combination with buforin II, checkerboard assays were conducted under the same conditions as the MICs but in Luria-Bertani (LB). Doubling dilutions of the two components – first temporin L vs temporin B and subsequently temporin B/temporin L vs buforin II - were performed on two 96-well plates, one horizontally and one vertically. These were combined and bacteria were added as for the MIC. FIC is calculated as (MIC of compound A in combination with B / MIC of compound A alone) + (MIC of compound B in combination with A / MIC of compound B alone). MICs were determined on the same plates as the FICs to increase reproducibility. FIC values ≤ 0.5 would be considered strongly synergistic and, consistent with a recent re-evaluation of FIC which stresses the importance of also measuring the MIC in the same microarray plate, values of 0.5 - < 1 were weakly synergistic.^17^ EMRSA-15 (NCTC 13616) and all other strains have been sequenced, to allow linkage of resistance phenotypes to known genetic traits.

In vitro *Pharmacodynamic assay* – *in vitro* pharmacodynamic assays were performed with epidemic methicillin resistant *S. aureus* 15 (EMRSA-15) (NCTC 13616) cultured in Mueller Hinton Broth (MHB). Cation adjusted MHB (CA-MHB) was used when testing daptomycin due to its requirement for Ca^2+^ ions for activity. Bacteria were cultured overnight in 10 ml of MHB or CA-MHB at 37°C and diluted just prior to plate inoculation to an OD_600_ of 0.002. Stock solutions of temporin B, temporin L, pleurocidin, tobramycin or gentamicin were prepared in sterile MilliQ water at a concentration of 200x MIC. Daptomycin was prepared in methanol at a concentration of 2000x MIC and diluted with media to 200x MIC in the first well. A dilution series was made in the top row of a polypropylene 96-well plate from 200x MIC to 0.2x MIC in a volume of 100 μl, to which 100 μl of the bacterial suspension was added to have a total of 1×10^6^ log-phase colony forming units (CFU) in 200 μl. The first t=0 sample was taken <30 seconds after addition of bacteria to the plate with further samples taken at appropriate intervals thereafter. Peptide challenged bacteria were sampled every 20 minutes for 120 minutes due to rapid killing while tobramycin, gentamicin and daptomycin challenged bacteria were sampled every hour for 6 hours. 15 μl was removed from each well and diluted 1:1000 in phosphate buffered saline and plated onto MH agar or CA-MH agar plates. The plates were incubated at 37°C overnight for CFU counting. CFU data were log_10_ transformed, and the bacterial net growth rate was determined from the increase or decrease in CFU during the time of exposure to the peptides or antibiotics as the coefficient of a linear regression of log_10_ CFU as a function of time. The intercept of the regression was fixed by forcing the regression lines through the first CFU measurement (0 min) at a given antimicrobial concentration. The pharmacodynamic function according to Regoes *et al*^18^ describes the relationship between bacterial net growth rate *ψ* and the concentration of an antimicrobial (a):

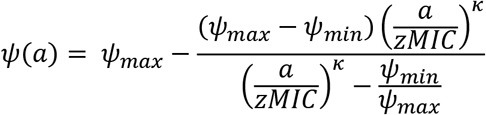

Fitting this function to the net bacterial growth rates in OriginPro 2020 (OriginLab Corporation, Northampton, MA) generates parameters *ψ_min_* and *ψ_max_*, respectively the minimum and maximum growth rate, zMIC, the pharmacodynamic minimum inhibitory concentration and κ, a measure of the cooperativity. Average parameters obtained from fits of three or more independently repeated experiments were compared by one-way ANOVA with Tukey post-hoc test. Since the CFU data is log_10_ transformed, the net growth rates, are thereafter reported to three significant figures

### Molecular dynamics simulations

Peptide starting structures were copies of the same conformer obtained from previous NMR calculations of peptide prepared in SDS micelles.^14,15^ Structural coordinates in the Protein Data Bank (www.rcsb.org) are accessed using codes 6GS5 and 6GIL for temporin L and temporin B respectively. Simulations were carried out on either the ARCHER Cray XC30 supercomputer or Dell Precision quad core T3400 or T3500 workstations equipped with a 1 kW Power supply (PSU) and two NVIDA PNY GeForce GTX570 or GTX580 graphics cards using Gromacs.^19^ The CHARMM36 all-atom force field was used in all simulations.^20,21^ The initial bilayer configuration was built using CHARMM-GUI.^22^ All membranes in this project contained a total of 512 lipids, composed of 1-palmitoyl-2-oleoyl-*sn*-glycero-3-phospho-(1’-*rac*-glycerol) (POPG) to reflect the lipid charge ratios of the plasma membrane of Gram-positive bacteria.^23,24^ Eight peptides were inserted at least 30 Å above the lipid bilayer in a random position and orientation at least 20 Å apart. The system was solvated with TIP3P water and Na^+^ ions were added to neutralize the total charge of the simulated system. Energy minimization was carried out using the steepest descent algorithm until the maximum force was less than 1000.0 kJ/ml/nm (^~^3000-4000 steps). Equilibration was carried out using the NVT ensemble for 100 ps and then a semi-isotropic NPT ensemble for 1000 ps with position restraints on the peptides. Hydrogen-containing bond angles were constrained with the LINCS algorithm. The production simulations were run using a semi-isotropic NPT ensemble using 2 femtosecond timesteps, with trajectories recorded every 2 picoseconds. All simulations were performed at a temperature of 310 K, which was controlled with a Nose-Hoover thermostat, and at a pressure of 1 bar, which was controlled with a Parrinello-Rahman barostat. All production simulations were run for a total of 200 ns and duplicated, with peptides inserted at different positions and orientations, giving a total of approximately 1.2 μs of simulation. To investigate the aggregation of the AMPs we have identified peptides that have come within 6 Å of each other at any given time step to be clustered. The connected components algorithm of NetworkX was used to find connectivity using graph theory. To quantify the conformation of the peptides, we measure torsion angles which are circular quantities, and the circular mean of psi or phi angles may be calculated as follows:

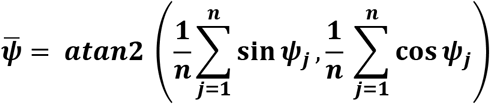

Similarly, the associated circular variance for psi or phi angles is calculated as follows:

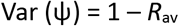

with *R* being given by:

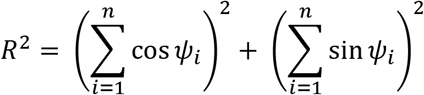

### Electrophysiology experiments (Patch-clamp)

Giant unilamellar vesicles (GUVs) composed of DPhPG were prepared in the presence of 1 M sorbitol by the electroformation method in an indium-tin oxide (ITO) coated glass chamber connected to the Nanion Vesicle Prep Pro setup (Nanion Technologies GmbH, Munich, Germany) using a 3-V peak-to-peak AC voltage at a frequency of 5 Hz for 140 minutes at 37°C.^25–27^ Bilayers were formed by adding the GUVs solution to a buffer containing 250 mM KCl, 50 mM MgCl_2_ and 10 mM Hepes (pH 7.00) onto an aperture in a borosilicate chip (Port-a-Patch^®^; Nanion Technologies) and applying 70-90 mbar negative pressure resulting in a solvent-free membrane with a resistance in the GΩ range. Diphytanoyl chains are used here for practical reasons since, unlike lipids with mixed palmitoyl-oleoyl chains such as POPG, these lipids do not undergo the main, temperature dependent transition from disordered fluid into the all *trans* configuration and remain in the same phase between −120° and +120°C ^28^ while, crucially, the membranes composed of these lipids are mechanically stable and have high specific resistance,^29^ essential for electrophysiology experiments. After formation, a small amount of peptide stock solution (in water) was added to 50 μL of buffer solution in order to obtain its active concentration. All the experiments were carried on with a positive holding potential of 50 mV. The active concentration, the concentration at which the peptide first showed membrane activity, for each peptide was obtained through a titration performed in the same conditions. For all the experiments a minimum of 6 concordant repeats were done. Current traces were recorded at a sampling rate of 50 kHz using an EPC-10 amplifier from HEKA Elektronik (Lambrecht, Germany). The system was computer controlled by the PatchControl™ software (Nanion) and GePulse (Michael Pusch, Genoa, Italy, http://www.ge.cnr.it/ICB/conti_moran_pusch/programs-pusch/software-mik.htm). The data were filtered using the built-in Bessel filter of the EPC-10 at a cut-off frequency of 10 kHz. The experiments were performed at room temperature. Data analysis was performed with the pClamp 10 software package (Axon Instruments). Estimation of pore radii was performed as previously.^30^

## Results

### Temporin B does not substantially increase the antibacterial potency of temporin L

FICs for the combination of temporin B and temporin L have been shown previously to be in the range of 0.55 to 0.75 for four Gram-positive strains and from 0.41 to 0.50 for four Gram-negative strains with a conservative value of ≤ 0.50 considered to represent synergy due to the inherent uncertainty in broth-microdilution assays.^12^ More recently, some researchers have suggested that values in the range 0.50 - 0.99 can also represent synergy - albeit modestly so - if care is taken to obtain MICs and FICs from the same plate.^17^ Values between 1.00 and 1.99 represent no interaction. In our previous studies, temporin L was shown to be more potent than temporin B against all strains included in both the Gram-positive and Gram-negative bacteria panels and *Candida albicans* (Table 1).^14,15^ Here, to facilitate a rapid and efficient screen of synergy in the whole panel, rather than employing checkerboard assays, three fixed ratios of temporin L and temporin B are tested to generate a range of FIC for three different stoichiometries (Table 1). In general, no evidence of strong synergy is found, with only small reductions of the amount of temporin L needed to inhibit bacterial growth when used in combination with temporin B. In some cases, a reduction in the amount of temporin L required is obtained with the addition of a small amount of temporin B and, considering its low potency when used alone, this produced FICs below 1.00. Overall, however, the present and previous studies agree that, at best, only modest synergistic improvements in potency are obtained by combining temporins L and B.

**Table 1.**
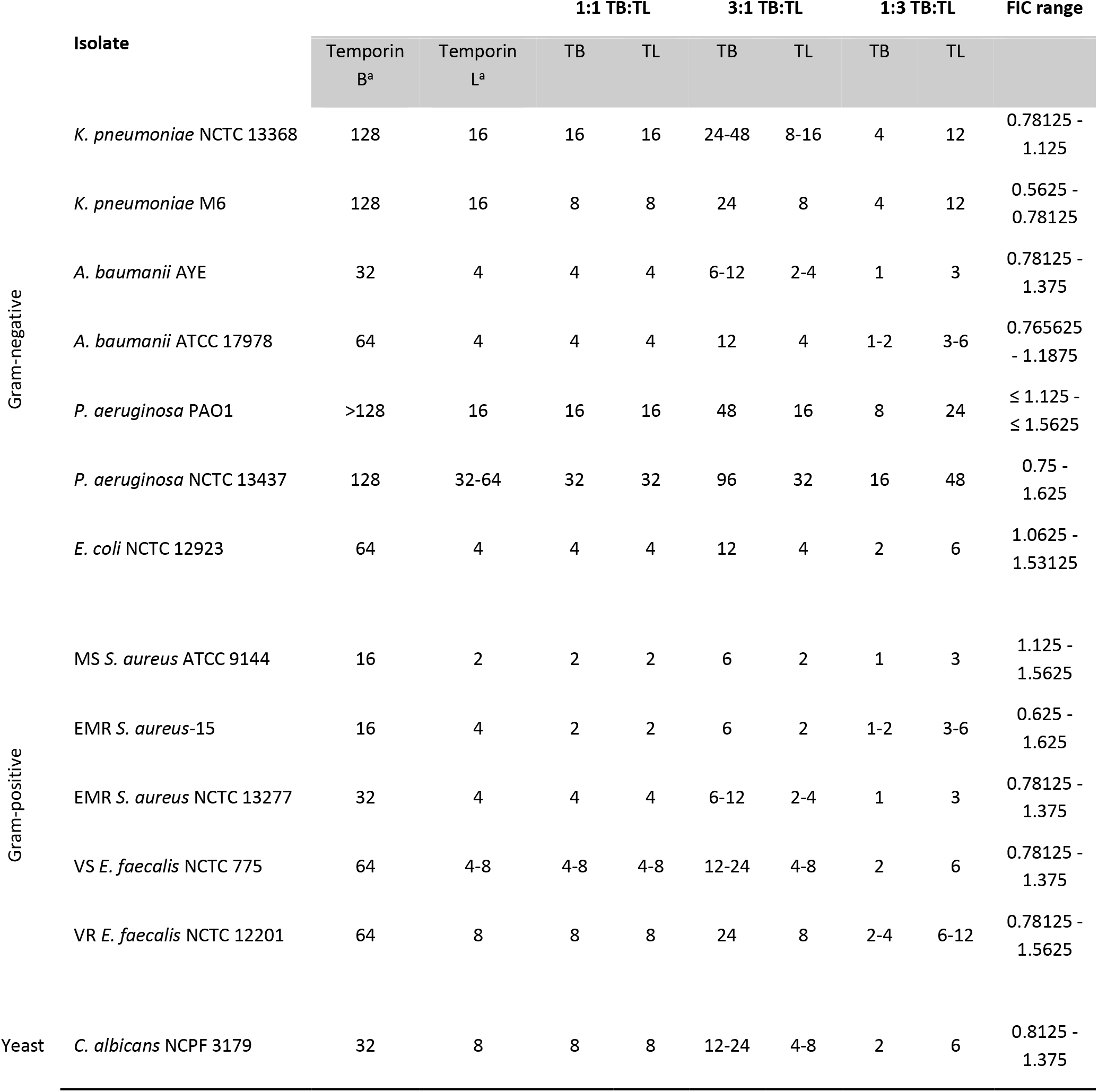
Antimicrobial activity. Data obtained from broth-microdilution assay in MHB. MS – methicillin sensitive; EMR – epidemic methicillin resistant; VS – vancomycin sensitive; VR – vancomycin resistant. MICs are reported in μg/ml. ^a^ data previously reported for temporin B^14^ and temporin L^15^. The MICs for temporin B and temporin L are given when used alone or in three combinations with differing stoichiometric ratios. The FIC range is the range of FICs obtained across the three differing stoichiometric ratios.

### Temporin B enhances the *in vitro* pharmacodynamic profile of temporin L when killing EMRSA-15

Of the panel strains where modest synergy is observed, EMRSA-15 is the most susceptible to both temporin L and temporin B. Here we determine the concentration dependent reduction in viable bacteria when log phase EMRSA-15 are challenged (Fig. 1) and present a comparison of the pharmacodynamic parameters obtained from challenges with AMPs – temporin L, a combination of temporin B and temporin L or, for comparison, pleurocidin^31^ - and existing clinically relevant antibiotics (Table 2). The bacteria were not challenged with temporin B alone since this AMP lacks potency and the synergy screen data (Table 1) indicate that, where modest synergy in potency is observed, the activity is largely attributed to temporin L, which is never present at less than ½ its MIC. While a variety of antibiotics are used to treat *Staphylococcus aureus* infections, many strains are now multi-drug resistant, only some antibiotics are bactericidal, and some may be restricted according to infection setting. EMRSA-15 is resistant to beta-lactams, second generation fluoroquinolones such as ciprofloxacin and third generation cephalosporins such as ceftazidime. It is sensitive to aminoglycosides including tobramycin and gentamicin, glycopeptides such as telavancin and vancomycin and daptomycin. All these may be bactericidal, but vancomycin has been found to have only bacteriostatic activity against some MRSA^32,33^ while use of daptomycin is more limited e.g. since its inhibition by pulmonary surfactant ensured it failed to meet noninferiority criteria in clinical trials for community-acquired pneumonia.

**Figure 1.**
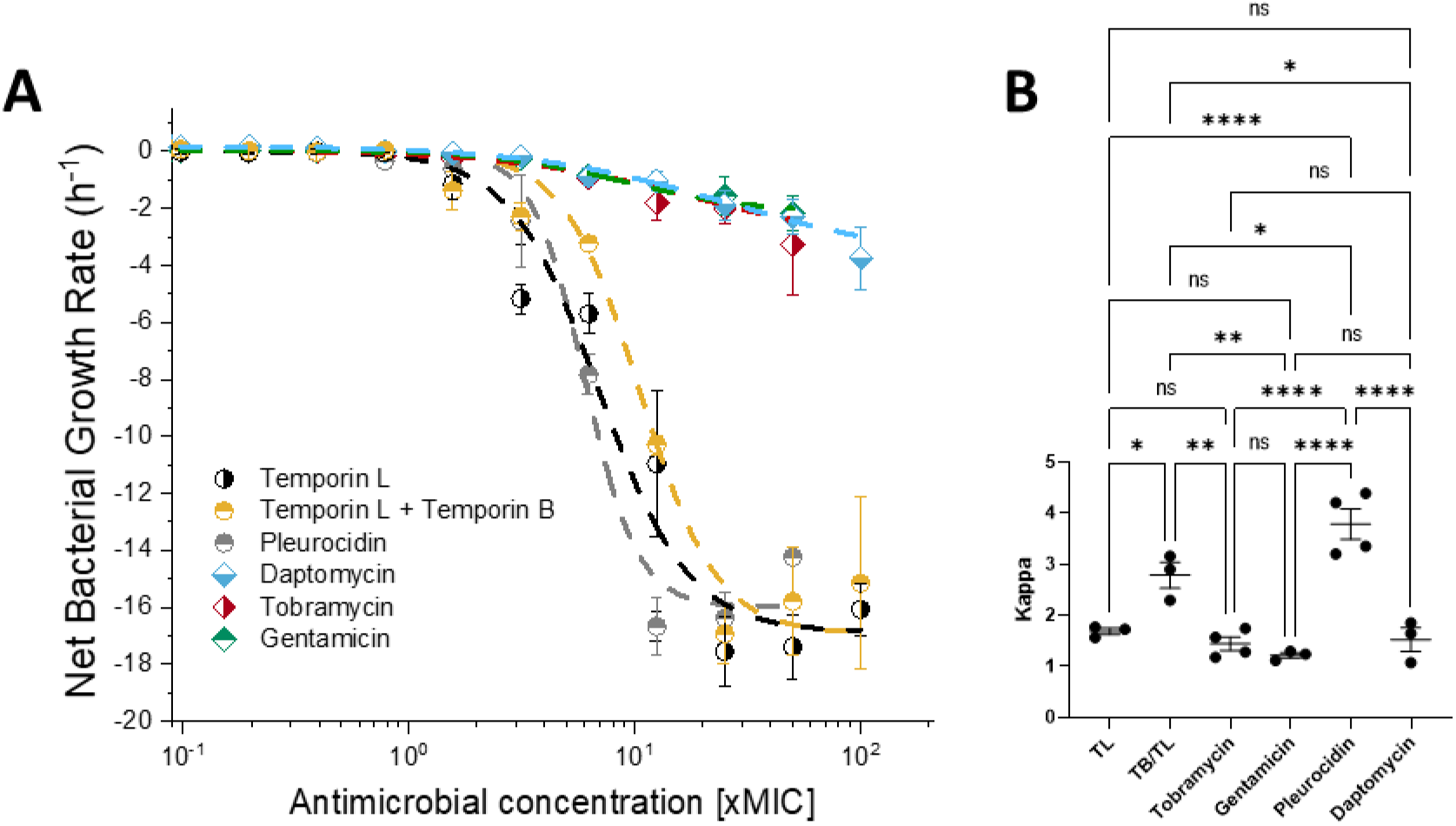
Pharmacodynamic response of EMRSA-15 to antibiotic challenge in MHB. EMRSA-15 was challenged with increasing concentrations of temporin L (TL), a 1:1 mol:mol ratio combination of temporin B and temporin L (TB:TL) or clinically relevant antibiotics. Curves shown are fits of averages of three independent repeated experiments (**A**). The cooperativity (kappa), pharmacodynamic MIC (zMIC) and maximum (*ψ*_max_) and minimum (*ψ*_min_) growth rates are provided in Table 2 while One-way ANOVA with Tukey post-hoc test multiple comparisons for kappa, highlight the differences in cooperativity between the AMPs and antibiotics (**B**). ns *p* > 0.05; * *p* < 0.05; ** *p* <0.01; *** *P* < 0.001; **** *p* < 0.0001

**Table 2.** Pharmacodynamic parameters obtained from challenge of EMRSA-15 in MHB with the indicated antibiotics. Parameters are the average and standard error of fits of three or more independently repeated experiments. * Assay conducted in CAMHB due to the requirement of daptomycin for Ca^2+^ ions for activity. Values that differ significantly (One-way ANOVA with Tukey post-hoc test; *p* < 0.01) with respect to temporin L are shown in bold.

| Condition | Kappa | zMIC (xMIC) | $\psi_{max}$ ($\text{h}^{-1}$) | $\psi_{min}$ ($\text{h}^{-1}$) |
| --- | --- | --- | --- | --- |
| Temporin L | $1.69 \pm 0.06$ | $0.30 \pm 0.08$ | $0.08 \pm 0.02$ | $-18.0 \pm 1.0$ |
| Temporin L / Temporin B | **$2.79 \pm 0.26$** | $0.42 \pm 0.16$ | $0.07 \pm 0.02$ | $-16.7 \pm 1.5$ |
| Pleurocidin | **$3.79 \pm 0.30$** | $1.18 \pm 0.10$ | $0.05 \pm 0.02$ | $-16.9 \pm 0.4$ |
| Tobramycin | $1.44 \pm 0.13$ | $0.64 \pm 0.16$ | $0.06 \pm 0.06$ | **$-3.36 \pm 0.83$** |
| Gentamicin | $1.22 \pm 0.05$ | $0.38 \pm 0.12$ | $0.08 \pm 0.04$ | **$-2.16 \pm 0.31$** |
| Daptomycin* | $1.52 \pm 0.23$ | $1.33 \pm 0.60$ | $0.07 \pm 0.04$ | **$-3.74 \pm 1.11$** |

Here linezolid, as expected, and vancomycin, are found to be bacteriostatic against EMRSA-15 and are not considered further. The peptides, and the clinically relevant daptomycin and aminoglycoside antibiotics tobramycin and gentamicin are bactericidal. However, pleurocidin, the combination of temporin L and temporin B, and temporin L alone all kill EMRSA-15 at a much faster rate than either aminoglycoside or daptomycin (*p* < 0.0001), as evidenced by *ψ_min_*, the minimum growth rate (Table 2). The cooperativity of the dose dependent activity, as characterised by the steepness of the slope in a dose-response curve and the parameter Kappa, reveals a potential benefit of challenging EMRSA-15 with a combination of temporin L and temporin B rather than temporin L alone. The cooperativity of the response to challenge with temporin L compares poorly with that to pleurocidin (*p* < 0.0001) (Table 2; Fig. 1B). However, when used in combination with temporin B, Kappa increases for the combination when compared with temporin L alone (*p* = 0.0315) (Fig. 1B, Table 2). Indeed, only the cooperativity of the response to challenge with the combination, but not temporin L alone, is greater than that achieved with either tobramycin (*p* = 0.0043) or gentamicin (*p* = 0.002).

The cooperativity of the dose dependent activity for both temporin L and the combination of temporin L/B is greater when the experiment is repeated in Luria-Bertani broth (Fig. S1). While the other parameters are similar in both media, in LB Kappa for the combination is nearly double that obtained in MHB. While in this media the addition of temporin B to temporin L alone does not increase Kappa (*p* = 0.9994), it does when buforin II is also present (*p* = 0.0023); LB is the only media in which we have found antibacterial activity with buforin II in broth microdilution assays,^34^ and here we identified synergy between buforin II and both temporin L (FIC = 0.56) and the combination of temporin L and temporin B (FIC = 0.5), but not temporin B (FIC = 1) using checkerboard assays. In contrast, adding buforin II to temporin L alone does not lead to any increase in Kappa (*p* = 0.2586) and may reduce it. The combination of all three peptides in LB produces a dose-response curve with a Kappa value almost six times that for the corresponding experiment with temporin L alone in MHB indicating there is substantial scope for the cooperativity of AMP bactericidal activity to vary according to the chemical environment.

### Temporin L and temporin B form hetero-oligomers in models of the Gram-positive plasma membrane

Since the *in vitro* pharmacodynamic study implies a possible interaction between temporin L and temporin B and since it is widely accepted that the main factor affecting the activity of AMPs is their interaction with the bacterial plasma cell membrane, we sought to identify whether either of these peptides modifies the membrane interaction of the other using first, all atom molecular dynamics simulations. We extend previous simulations of either eight temporin B^14^ or eight temporin L^15^ peptides binding to a 512 POPG lipid bilayer from 100 to 200 ns and perform new duplicate, 200 ns simulations of 4:4 combinations of temporin B and temporin L binding to the same bilayer. This allows us to assess the effect of temporin L and temporin B interaction on the peptide conformation and its flexibility (Fig. S2, S3); binding and insertion (Fig. S4); peptide-lipid hydrogen bonding (Fig. S5; S6); the formation of both homo- and hetero-oligomers in the bilayer (Fig. 2); and peptide induced lipid disordering (Fig. 3).

**Figure 2.**
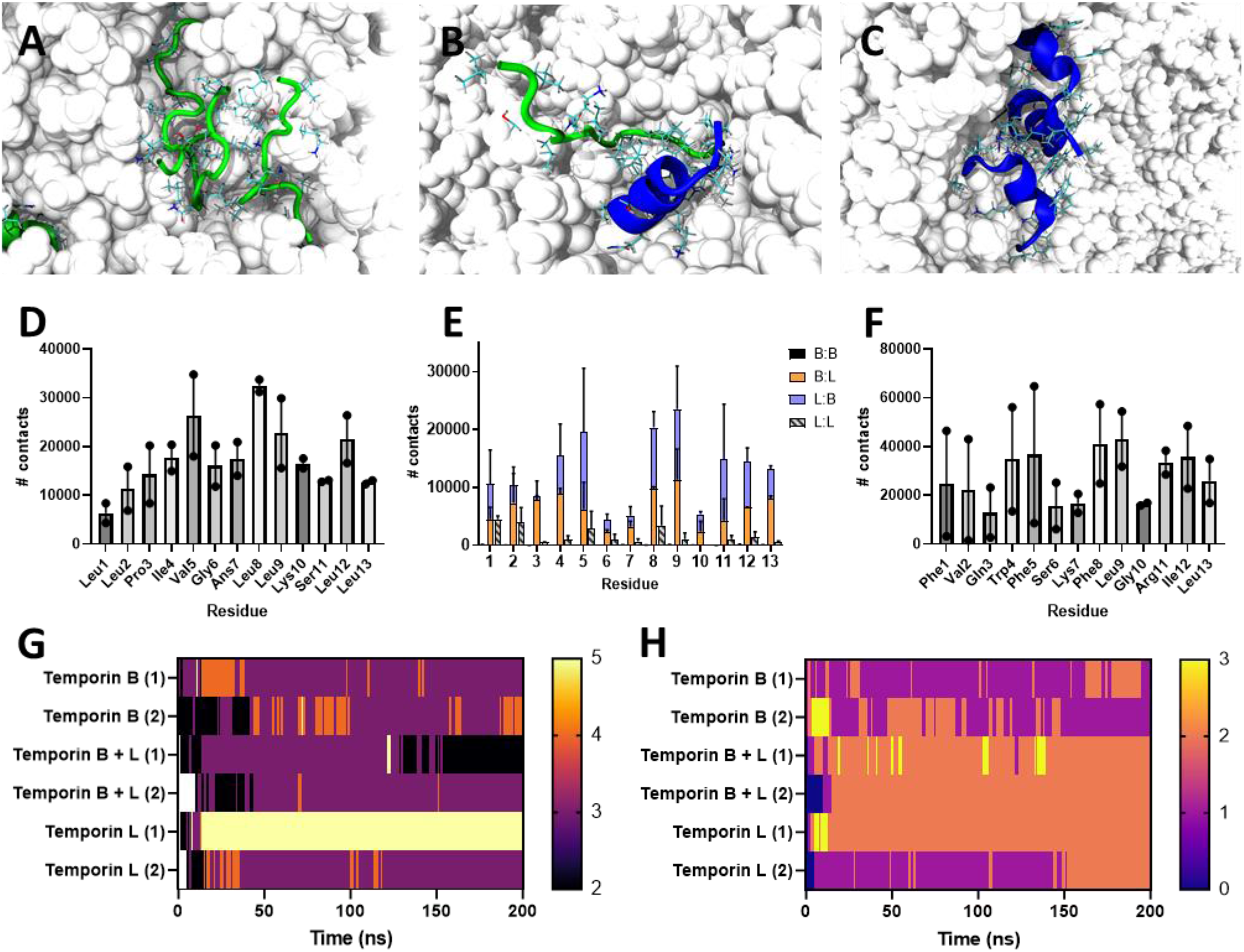
Temporin L and temporin B form hetero-oligomers in MD simulations of POPG bilayer challenge. Top zoom views of snapshots (A-C) and analysis of the average number of contacts for each residue involved in any homo- or hetero-oligomerisation (D-F) in simulations of eight temporin B (A/D), four temporin L (blue) and four temporin B (green) (B/E) or eight temporin L (C/F) peptides inserting into a 512 POPG lipid bilayer. Time-resolved analysis of the maximum number of peptides in any assembly (G) and the number of any such assemblies (H).

**Figure 3.**
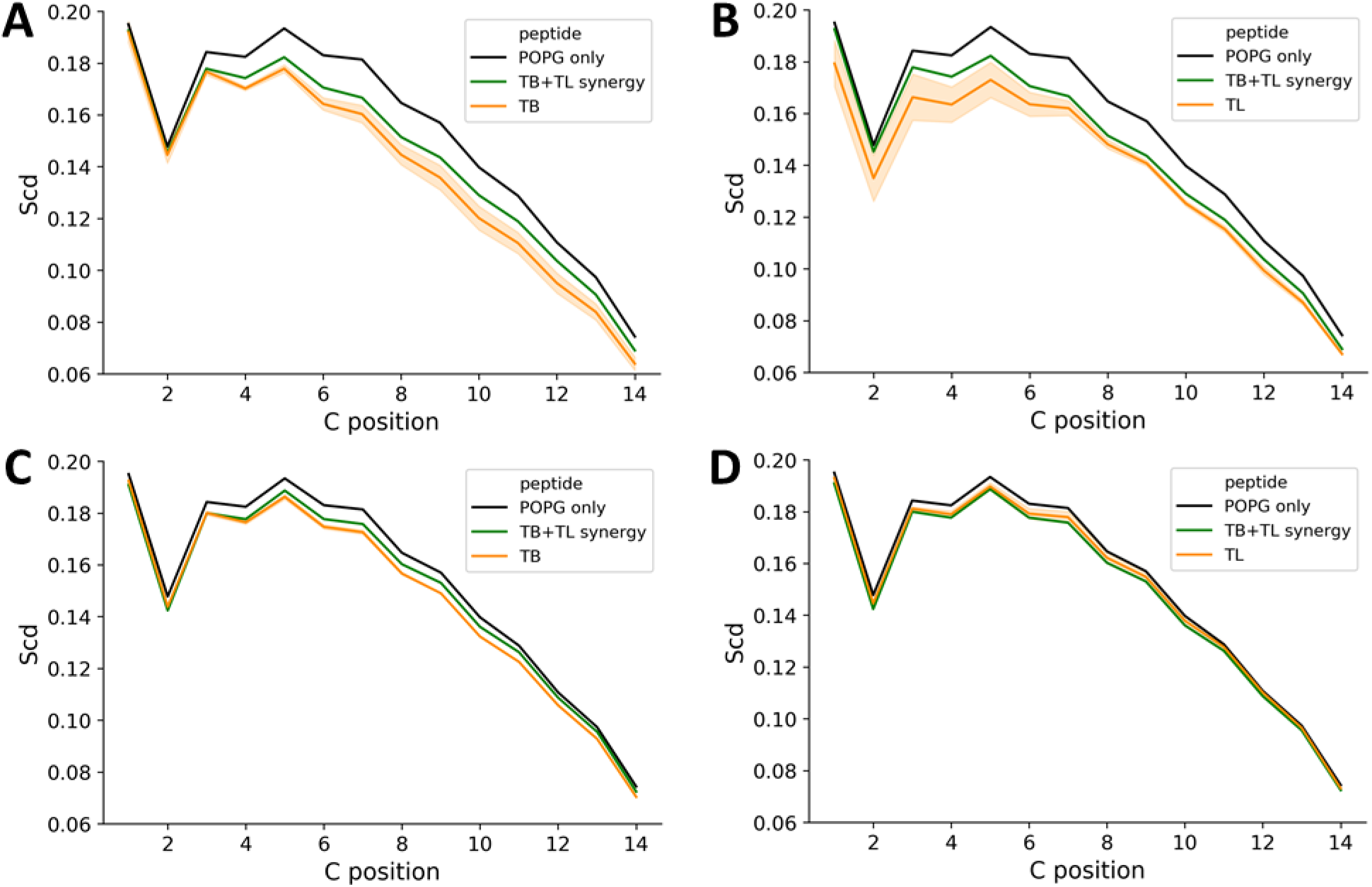
Hetero-oligomerisation reduces local membrane disordering by temporin L in MD simulations of POPG bilayer challenge. Order parameter profiles, averaged over the duration of the 200 ns simulations, are shown for lipids within 4 Angstroms of each inserting peptide (A, B) or for the whole bilayer (C, D). Comparisons are provided for temporin B (A, C) or temporin L (B, D). Data is an average of two independently repeated simulations for each condition.

On binding to the POPG bilayer, temporin B does not adopt α-helix conformations, as the Ramachandran contour plots, representing phi and psi dihedral angles averaged over the duration of the simulation, indicate that the peptides adopt a type II β-turn conformation (Fig. 2SA). In contrast, the majority of residues in temporin L (Trp4-Gly10) do adopt α-helix conformation. However, some residues may also adopt the type II β-turn (Fig. S2C). These residues are located at the N- and C-termini and these regions also exhibit greater conformational flexibility, as measured by the circular variance of the psi dihedral angle (Fig. S3G). When temporin L is combined with temporin B, conformational flexibility in temporin L is reduced across the whole peptide (Fig. S3C/D), in particular in the N- and C-termini (Fig. S3G/H), the α-helix conformation is extended (Gln3-Arg11) and evidence of type II β-turn diminishes (Fig. S2D). In contrast, temporin B experiences considerable conformational flexibility whether temporin L is present or not (Fig. S3A/B/E/F) and there are no notable changes in conformation (Fig. S2B).

Since previous work has shown that temporin L acts to prevent oligomerisation of either temporin A or temporin B in lipopolysaccharide (LPS), and we have separately shown that both temporin B and temporin L form oligomers when they insert into model bilayers^14,15^ we next assessed whether either peptide inhibited the membrane penetration (Fig. S3), the interaction of the peptides with the bilayer (Fig. S5; S6) and characterised any aggregates that formed (Fig. 2). As previously, both temporin B and temporin L penetrate the membrane via their N-termini and this is not substantially altered when the two different peptides are applied to the membrane in combination (Fig. S4).

The initial insertion is completed within 75 ns in all simulations, though penetration of temporin B is a little faster and deeper in the combination than when applied alone (Fig. S4A). Penetration of either temporin B or L is therefore not inhibited by the presence of the other temporin and, indeed, that of temporin B may be facilitated by temporin L. Consistent with this, neither the total number of peptide-lipid hydrogen bonds (Fig. S5) nor the pattern of residue specific peptide-lipid hydrogen bonds (Fig. S6) is altered, for either temporin L or temporin B, when membranes are challenged with the peptides in combination.

The effect of combining temporin B and temporin L is clearer following analysis of peptide-peptide oligomerisation in the membrane (Fig. 2). As can be observed (Fig. 2B), and as was predominantly the case throughout the 200 ns duration of the simulation (Fig. 2G), small hetero-oligomers, usually comprising two or three peptides, formed in POPG membranes. For temporin B, trimers (Fig. 2A) and occasionally tetramers were frequently observed in analogous simulations,^14^ and these are now shown to endure throughout the extended simulation (Fig. 2G), while temporin L forms trimers (Fig. 2C), tetramers and, in one simulation, a stable pentamer,^15^ and these are retained as the simulation is extended to 200 ns (Fig. 2G). For the combination there are only four, as opposed to eight, of each temporin molecule in each simulation and this may impact the probability of higher or lower order oligomerisation and the ability to draw conclusions about the size distribution of resulting pores. However, the likely preference of each peptide for hetero-oligomerisation over homo-oligomerisation does provide support for a synergistic effect in the target membrane. This is revealed by analysis of the number of contacts between peptide monomers in each simulation (Fig. 2D-F). By chance, hetero-oligomeric contacts should predominate over homo-oligomeric contacts at a ratio of 4:3. Instead homo-oligomers of either temporin B or temporin L are very rare while hetero-oligomers are much more frequent (hetero- to homo-ratios 158:1 temporin B; 5.4:1 temporin L). With the exception of Arg11 in temporin L, the residues in each peptide involved in mediating assembly are hydrophobic, are located in the same positions in both temporin B and L, do not change substantially whether hetero- or homooligomers are being formed and are not involved in hydrogen bonding with the lipid bilayer (Fig. 2D-F; S6).

While spontaneous pore formation in membranes is rarely observed in simulations when peptides start in the water phase,^35^ the lipid disorder associated with their formation, in such studies,^36,37^ is observed for some peptides irrespective of whether pores form or not.^14,38,39^ The lipid disorder is greatest in those lipids associated with the peptide while order may increase for non-associated lipids,^36,37^ and in our previous work the same effect was observed for magainin 2, pleurocidin and its analogues, and temporin B.^14,31,38^ Here, both temporin B (Fig. 3A) and temporin L (Fig. 3B) are observed to strongly disorder POPG lipids located within 4 Å of any peptide, with temporin L having a greater effect. In contrast, the disordering effect of temporin L on the whole bilayer is less noticeable compared with that of temporin B (Fig. 3C/D). When the peptides are applied in combination, the local disordering effect of both peptides is attenuated (Fig. 3A/B) while the impact on the whole bilayer is intermediate between that achieved with either peptide alone (Fig. 3C/D).

### Temporin B modulates channel activity induced by temporin L in model membranes

We made use of the Port-a-patch^®^ automated patch-clamp system from Nanion Technologies (Munich, Germany) to determine whether the addition of temporin B modifies the ability of temporin L to disrupt DPhPG bilayers, mimicking Gram-positive bacteria cytoplasmic membranes (Fig. 4).^23,24^ Our experimental approach involves finding the lowest concentration of peptide that induces detectable conductance and then measuring the latency – the time taken for conductance to commence after addition of peptide -, recording whether the membrane is ultimately broken and quantifying any characteristic channel-like activity – well-defined events with discrete opening levels. Previously, we showed that temporin B does induce conductance in DPhPG bilayers but at a relatively high concentration of 35 μM.^14^ It induces irregular conductance activity, and no evidence of regular channel formation was detected. Conductance activity does however appear relatively quickly after temporin B administration, and the membrane soon ruptures. In contrast, we have previously shown that temporin L does induce channel like activity that endures, and this is achieved with less peptide (10 μM) than is required for temporin B.^15^ Here, we find that combining temporin B and temporin L, in a 3.5:1 molar ratio reflecting their differing potency when used alone, substantially affects the ability to induce conductance. In combination, the concentrations of the peptides required to induce conductance are twelve-fold lower than when each peptide were used alone. Channel-like activity is detected (Fig. 4A) and it appears more rapidly than when temporin L is applied alone (Fig. 4C). However, the amplitude, conductance and estimated pore radii are much smaller than those observed for temporin L alone (Fig. 4A/B, Table 3). Patch-clamp therefore reveals the combination of temporin L and temporin B induces channel-like activity at much lower concentrations and faster, but the channels are much smaller than achieved with temporin L alone.

**Figure 4.**
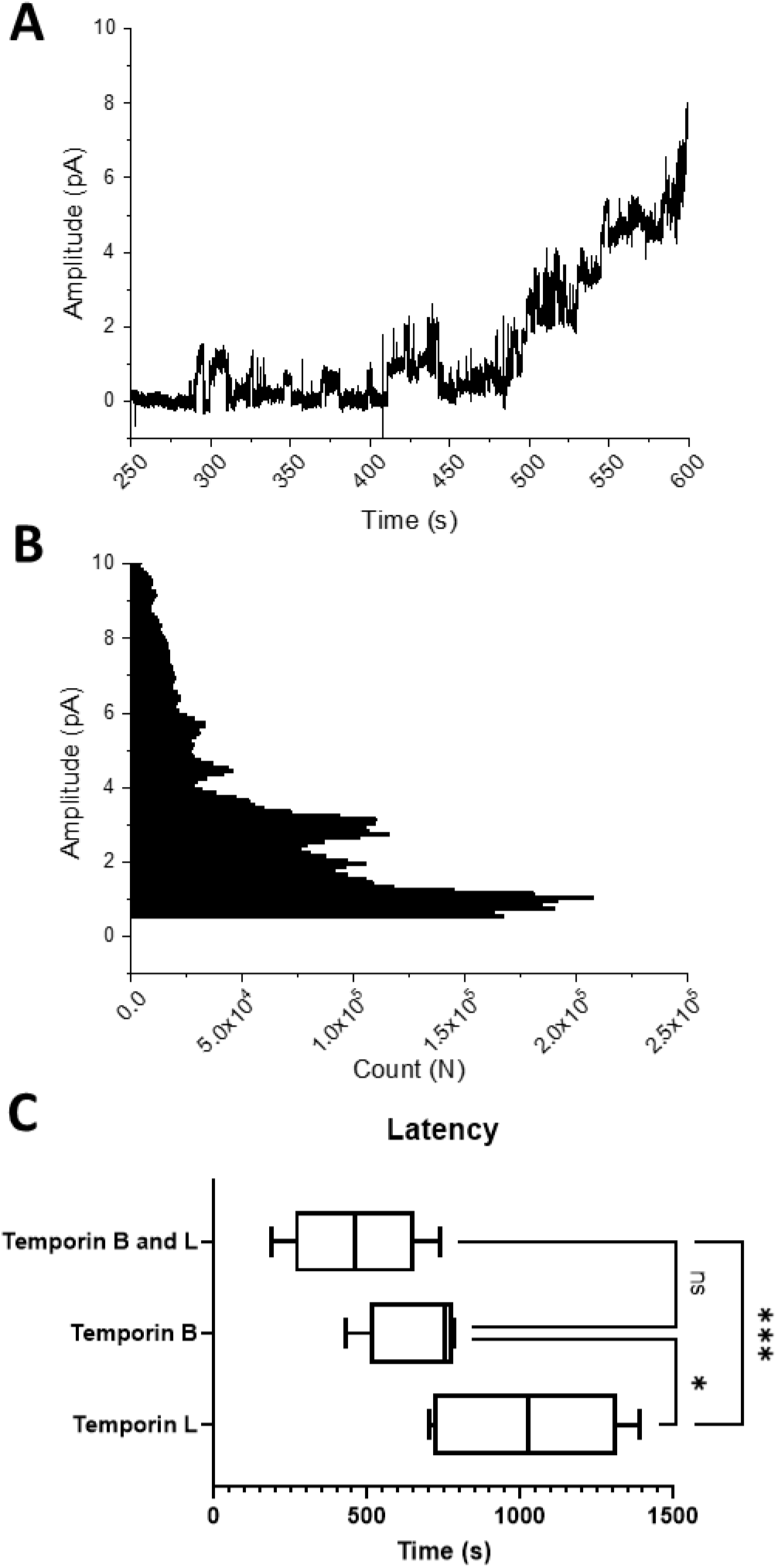
Patch-clamp analysis of challenge of a DPhPG bilayer with a combination of temporin L and temporin B. The concentrations, of temporin L (0.84 μM) and temporin B (2.92 μM) used, correspond to the minimum amount of the combination needed to induce conductance and are equal to 1/12 of the concentrations needed to induce conductance when each peptide is applied alone. A representative of six traces (A) together with a frequency plot of events of varying amplitude across all six traces (B). The average time taken for conductance to begin after peptide addition (latency) shows conductance begins more rapidly for the combination than temporin L alone (C); One-way ANOVA with Tukey post-hoc test, * *p* < 0.05, *** *p* < 0.001.

**Table 3.** Summary of channel-like activity detected at various opening levels. DPhPG membranes were challenged with 10 μM temporin L alone or a combination of 0.84 μM temporin L and 2.92 μM temporin B. Temporin B alone does not induce channel-like activity. Level 1 is present in 5/6 traces acquired, Level 2 is present in 3/6 out of 6 traces and Level 3 is present in 2/6 traces. * data previously reported^15^.

| Peptide | Parameter | Parameter |  |  |
| --- | --- | --- | --- | --- |
|  |  | Level 1 | Level 2 | Level 3 |
| Temporin L * | Amplitude (pA) | $0.89 \pm 0.02$ | $25.4 \pm 0.1$ | - |
| Temporin L / Temporin B | | $0.42 \pm 0.03$ | $2.17 \pm 0.33$ | $4.77 \pm 0.58$ |
| Temporin L * | Conductance (pS) | $17.9 \pm 0.4$ | $507 \pm 2$ | - |
| Temporin L / Temporin B | | $8.40 \pm 0.60$ | $43.4 \pm 6.6$ | $95.4 \pm 11.6$ |
| Temporin L * | Estimated pore radius (nm) | $0.08 \pm 0.01$ | $0.43 \pm 0.03$ | - |
| Temporin L / Temporin B | | $0.05 \pm 0.01$ | $0.12 \pm 0.05$ | $0.18 \pm 0.06$ |

## Discussion

The discovery of the temporin peptides^11^ in the skin secretion of *Rana temporaria* has precipitated a large body of work seeking to understand and develop those AMPs with the greatest antimicrobial activity into useful antibiotics.^9,10,39^ Ten temporin peptides were initially described and they share extensive sequence similarity, ranging from 76.9% to 100% relative to temporin B (Table 4). Though all were active against *Bacillus megaterium*, only temporins A, B, F, G and L were active against *Escherichia coli* when tested individually i.e. those carrying at least a +2 nominal charge and 13 residues in length. Subsequently, attention has been largely focused on temporins L, B and A,^39–50^ despite temporins F and G being produced at similar levels to temporins A and B, and temporin C being the most abundant of them all.^11^ Temporin L has greater antibacterial potency against Gram-negative bacteria, binds lipopolysaccharide and hence has antiendotoxin properties but is also relatively cytotoxic.^40–47^ Temporins A and B are more active against Gram-positive bacteria though analogues of temporin B have been produced with a broader spectrum of activity.^49^ Temporins A and B have also been shown to act in synergy with temporin L against Gram-negative bacteria and the mechanism for this synergy has been explored using biophysical methods.^12,50^ However, since the other temporins have received less attention, it is unclear what the biological benefits are of producing such a set of closely related peptides nor what the relatively minor changes in amino acid sequence, at least between temporins A-K (Table 4), achieve.

**Table 4.** Alignment of temporin peptides sequences and their physicochemical characteristics. Average hydrophobicity is given on the whole-residue hydrophobicity octanol-interface scale (Δ*G*_woct_ - Δ*G*_wif_) based on the free energy of transfer from water to palmitoyloleoylphosphatidylcholine and to *n*-octanol.^60^ All peptides are considered amidated at the C-terminus, but this is not considered in the hydrophobicity calculation. In temporins A-G, Pro3, Gly6, Leu9 and Leu13 are absolutely conserved. Leu9 and Leu13 are also conserved in temporin L.

| Peptide | Sequence | Charge | aa | $\Delta G_{\text{woct}} - \Delta G_{\text{wif}}$<br>pH 8 |
| --- | --- | --- | --- | --- |
| Temporin A | FLP LIG**R**VLSGIL-NH<sub>2</sub> | +2 | 13 | -0.169 |
| Temporin B | LLP IVGNLL**K**SLL-NH<sub>2</sub> | +2 | 13 | -0.160 |
| Temporin C | LLP ILGNLLNGLL-NH<sub>2</sub> | +1 | 13 | -0.216 |
| Temporin D | LLP IVGNLLNSLL-NH<sub>2</sub> | +1 | 13 | -0.266 |
| Temporin E | VLP IIGNLLNSLL-NH<sub>2</sub> | +1 | 13 | -0.275 |
| Temporin F | FLP LIG**K**VLSGIL-NH<sub>2</sub> | +2 | 13 | -0.110 |
| Temporin G | FFP VIG**R**ILNGIL-NH<sub>2</sub> | +2 | 13 | -0.162 |
| Temporin H | LSP NLL**K**SLL-NH<sub>2</sub> | +2 | 10 | -0.086 |
| Temporin K | LLP NLL**K**SLL-NH<sub>2</sub> | +2 | 10 | -0.188 |
| Temporin L | FVQWFS**K** FLG**R**IL -NH<sub>2</sub> | +3 | 13 | -0.018 |

The previous studies of synergy between temporin L and either temporins A or B have focussed on Gram-negative bacteria to understand gains in antibacterial potency. These reveal that temporin L affects the oligomerisation of temporin B in lipopolysaccharide environments.^12,50^Here, though FIC data would indicate that there is no interaction between temporin L and temporin B save for possibly a weak synergistic effect in some strains (additive elsewhere), three separate pieces of evidence indicate there is an interaction between temporin L and temporin B that will also influence their combined activity against Gram-positive bacteria.

First, we show that the addition of temporin B to either temporin L in MHB or temporin L/buforin II in LB increases the cooperativity of the dose dependent rate of bacterial killing and that this is greater than that observed for antibiotics. A limitation of our study is the absence of *in vitro* PD data for temporin B alone, and we cannot conclude whether the combination of temporin L and temporin B has greater cooperativity than both individual components or whether temporin B has higher cooperativity than temporin L and the combination then matches this. Instead, since the MIC of temporin L in combination with temporin B is never less than half of its MIC when applied alone, we have demonstrated that the effect of the combination is to retain the potency of temporin L but with enhanced cooperativity derived from the addition of temporin B, well below its own MIC.

The effects and mechanisms of adding buforin II, as a component of antimicrobial peptide combinations, warrant further investigation, not least because the effect of its addition in the present study is not clear cut. The present observations are reported here for two reasons. First, buforin II is a twenty-one amino acid histone H2A fragment, initially identified in the Asian toad *Bufo bufo garagrizans*,^51^ but its sequence is also found in the *Rana temporaria* (and mammalian) histone H2A and hence there is potential for it interacting with a wide variety of AMPs in different organisms. Second, buforin II accumulates within bacteria, has high affinity for nucleic acids and its antibacterial mechanism of action is independent of membrane lysis and hence completely different to that of either temporin L or temporin B.^51–53^ Therefore, the increases in cooperativity, obtained by combining temporin B with temporin L alone (in MHB) or with buforin II (in LB), are two examples of AMPs with differing mechanisms from the same organism combining to produce bactericidal activity with greater cooperativity. This is consistent with previous work that has shown diverse AMPs, but from different organisms, display greater cooperativity than antibiotics when killing *Escherichia coli* MG1655 and that this is enhanced when these AMPs are used in three-way combinations.^7,54^

The present data indicate that the possible impact of bacterial growth conditions, and other factors, on AMP pharmacodynamics should be explored in more depth, but are sufficient to conclude that combining AMPs has potential in further distinguishing their *in vitro* pharmacodynamic properties from those of bactericidal antibiotics. By extension, future work may now test the theory that the more cooperative pharmacodynamic profile achieved with combinations of AMPs mitigates the risk of resistance developing and hence a rationale for the evolution of synergistic AMP families within individual species.

Second, we show that temporin B and temporin L readily form hetero-oligomers in MD simulations of challenge of a model of the Gram-positive plasma membrane. Although there are many ways in which two different AMPs may influence the activity of each other, the formation of hetero-oligomers has been observed for other AMPs that are known to act in synergy; magainin 2 and PGLa, which are structurally related and from the same organism (*Xenopus laevis*), is a very well-studied example.^55–59^ In coarse-grain MD simulations, magainin 2 was shown to fix the membrane inserting state of PGLa, which otherwise continuously inserts and leaves the membrane, and aid recruitment of other peptides into heterodimers involved in the formation of transmembrane pores,^59^ explaining the observed increase in membrane affinity of the mixture.^58^ Here we use atomistic simulations to sample a shorter timescale, but while the membrane insertion of either temporin L or temporin B is largely unaffected by the presence of the other, the observation of hetero-oligomer formation, and concomitant restriction of temporin L homo-oligomer formation and lipid acyl-chain disordering can be expected to be manifested in altered disruptive effects of the peptides on the target plasma membrane.

Third, we use conductance measurements to show that the interaction between the temporin L/temporin B combination and model membranes fundamentally differs to that observed when either peptide is applied alone with conductance events observed more quickly and with much lower amounts of each peptide when applied in combination than when applied alone. The conductance manifests as regular channel-like events, similar to those produced with temporin L alone, but of a much lower conductance and calculated size. To achieve greater cooperativity the combination should suppress bactericidal activity at lower AMP concentrations but enhance it at higher concentrations. It is possible that the ability to induce conductance in model membranes with much less peptide, and faster, is a manifestation of enhanced bactericidal membrane activity of the temporin B/temporin L combination. However, unless other factors intervene to substantially reduce antibacterial activity overall, we would also expect to see a considerable increase in antibacterial potency for the combination. However, this is inconsistent with the modest synergy observed, as described by the FIC. Alternatively, the low conductance events observed for the combination may be insufficient for a bactericidal effect and this then would be consistent with temporin B preferentially forming hetero-oligomers with temporin L that are less effective than temporin L homo-oligomers. Only at higher relative concentrations of temporin L do high conductance channels form and hence cooperativity is enhanced. Therefore, while the present biophysical data establishes high probability of an interaction between temporin L and temporin B in the target plasma membrane and provides clues as to how the greater cooperativity in bactericidal activity is achieved, a complete mechanistic understanding will require a future investigation of dose-dependent effects in both patch-clamp studies and MD simulations.

## Conclusion

Combining two or, potentially, more antimicrobial peptides from the same organism improves the *in vitro* pharmacodynamic properties of the bactericidal action. For temporin L and temporin B, this is likely achieved through modification of aggregates formed by the peptides in the target membrane and the resulting ability of temporin L to induce channel-like conductance and suggests an evolutionary benefit for generating a family of AMPs and a more important role for those AMPs that alone have low antibacterial potency.

## ASSOCIATED CONTENT

### Supporting Information

Further *in vitro* pharmacodynamic data and analysis of the MD simulations is provided as Supplementary Figures S1-S6.

## AUTHOR INFORMATION

### Author Contributions

GM, DAP and AJM designed the study. AJM wrote the main manuscript text and, together with PMF, MaC and GM, prepared all figures. MaC and GM designed and performed *in vitro* pharmacodynamic assays. GM performed and analysed patch-clamp measurements. PMF, MaC and CDL designed, performed and/or analysed atomistic simulation data. CKH, MeC, MaC and JMS conducted antimicrobial susceptibility testing. All authors approved the manuscript.

### Notes

The authors declare no competing interests.

## Acknowledgment

CDL acknowledges the stimulating research environment provided by the EPSRC Centre for Doctoral Training in Cross-Disciplinary Approaches to Non-Equilibrium Systems (CANES, EP/L015854/1). PMF was supported by a Health Schools Studentship funded by the EPSRC (EP/M50788X/1). This work used the ARCHER UK National Supercomputing Service (http://www.archer.ac.uk). Part of TOC graphic created with BioRender.com.

## Uniprot accession

Uniprot accession codes are as follows: Temporin L (P57104), temporin B (P79874), buforin II (P55897) and pleurocidin (P81941).

## For Table of Contents use Only

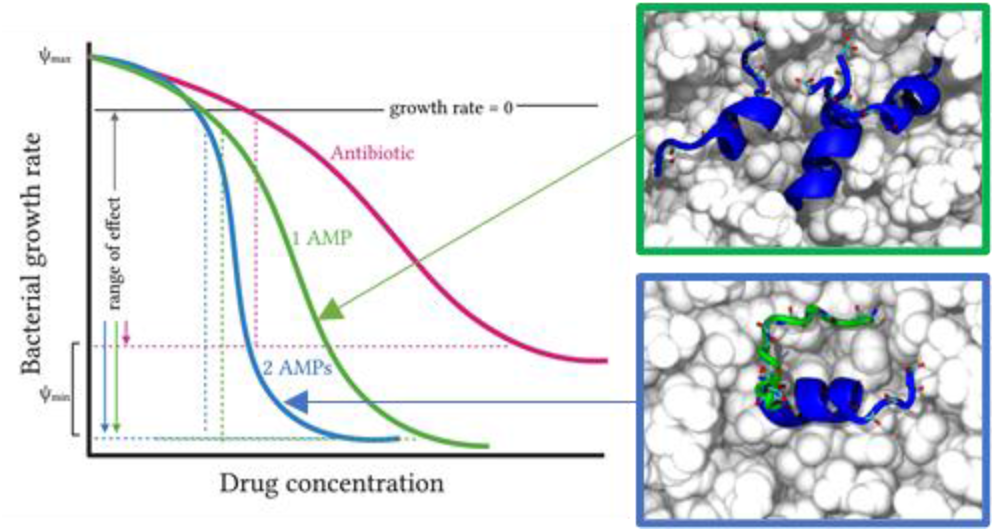

## Notes

### Competing Interest Statement

The authors have declared no competing interest.

## REFERENCES

1. Mookherjee, N., Anderson, M.A., Haagsman, H.P. & Davidson, D.J. Antimicrobial host defence peptides: functions and clinical potential. Nature Reviews Drug discovery 19, 311–332 (2020).

2. Jangir, P.K., Ogunlana, L. & MacLean, R.C. Evolutionary constraints on the acquisition of antimicrobial peptide resistance in bacterial pathogens. Trends in Microbiology. Doi: https://doi.org/10.1016/j.tim.2021.03.007 (2021)

3. Kubicek-Sutherland, J.Z., Lofton, H., Vestergaard, M., Hjort, K., Ingmer, H. & Andersson, D.I. Antimicrobial peptide exposure selects for *Staphylococcus aureus* resistance to human defence peptides. J. Antimicrob. Chemother. 72, 115–117 (2017).

4. Spohn, R., Daruka, L., Lázár, V., Martins, A., Vidovics, F., Grézal, G., Méhi, O., Kintses, B., Számel, M., Jangir, P.K., Csörgő, B., Györkei, A., Bódi, Z., Faragó, A., Bodai, L., Földesi, I., Kata, D., Maróti, G., Pap, B., Wirth, R., Papp, B. & Pál, C. Integrated evolutionary analysis reveals antimicrobial peptides with limited resistance. Nat. Commun. 10, 4538 (2019).

5. Yang, Q., Li, M., Spiller, O.B., Hinchliffe, P., Li, H., MacLean, C., Niumsup, P., Powell, L., Pritchard, M., Papkou, A. & Shen, Y. Balancing *mcr-1* expression and bacterial survival is a delicate equilibrium between essential cellular defence mechanisms. Nat. Commun. 8, 2054 (2017).

6. Lazzaro, B.P. et al. (2020) Antimicrobial peptides: Application informed by evolution. Science 368, eaau5480

7. Yu G., Baeder, D.Y., Regoes, R.R. & Rolff, J. Predicting drug resistance evolution: Insights from antimicrobial peptides and antibiotics. Proc. Biol. Sci. 285, 20172687 (2018).

8. Kintses, B., Jangir, P.K., Fekete, G., Számel, M., Méhi, O., Spohn, R., Daruka, L., Martins, A., Hosseinnia, A., Gagarinova, A., Kim, S., Phanse, S., Csörgő, B., Györkei, Á., Ari, E., Lázár, V., Nagy, I., Babu, M., Pál, C. & Papp, B. Chemical-genetic profiling reveals limited cross-resistance between antimicrobial peptides with different modes of action. Nat. Commun. 10, 5731 (2019).

9. Romero, S.M., Cardillo, A.B., Martínez Ceron, M.C., Camperi, S.A. & Giudicessi, S.L. Temporins: an approach of potential pharmaceutic candidates. Surgical Infections 21, 309–322 (2020).

10. Bhattacharjya, S. & Straus, S.K. Design, engineering and discovery of novel α-helical and β-boomerang antimicrobial peptides against drug resistant bacteria. Int. J. Mol. Sci. 21, 5773 (2020).

11. Simmaco, M. et al. Temporins, antimicrobial peptides from the European red frog *Rana temporaria*. Eur. J. Biochem. 242, 788–792 (1996).

12. Rosenfeld, Y., Barra, M., Simmaco, M., Shai, Y. & Mangoni, M.L. A synergism between temporins towards Gram-negative bacteria overcomes resistance imposed by the lipopolysaccharide protective layer. J. Biol. Chem. 281, 28565–28574 (2006).

13. Carotenuto, A. et al. A different molecular mechanism underlying antimicrobial and hemolytic actions of temporins A and L. J. Med. Chem. 51, 2354–2362 (2008).

14. Manzo, G., Ferguson, P.M., Gustilo, V.B., Hind, C., Clifford, M., Bui, T.T., Drake, A.F., Atkinson, R.A., Sutton, M.J., Batoni, G., Lorenz, C.D. Phoenix, D.A. & Mason, A.J. Minor sequence modifications in temporin B cause drastic changes in antibacterial potency and selectivity by fundamentally altering membrane activity. Scientific Reports 9, 1385 (2019).

15. Manzo, G, Ferguson, P.M., Hind, C., Clifford, M., Gustilo, V.B., Ali, H., Bansal, S.S., Bui, T.T., Drake, A.F., Atkinson, R.A., Sutton, J.M., Lorenz, C.D. Phoenix, D.A. & Mason, A.J. Temporin L and aurein 2.5 have identical conformations but subtly distinct membrane and antibacterial activities. Scientific Reports 9, 10934 (2019).

16. Wiegand, I., Hilpert, K., & Hancock, R.E. Agar and broth dilution methods to determine the minimal inhibitory concentration (MIC) of antimicrobial substances. Nature Protocols 3, 163–175 (2008).

17. Fratini F, Mancini S, Turchi B, Friscia E, Pistelli L, Giusti G & Cerri D. (2017) A novel interpretation of the Fractional Inhibitory Concentration Index: The case *Origanum vulgare* L. and *Leptospermum scoparium* J. R. et G. Forst essential oils against *Staphylococcus aureus* strains. Microbiol. Res. 195, 11–17.

18. Regoes, R.R., Wiuff, C., Zappala, R.M., Garner, K.N., Baquero, F. & Levin, B.R. Pharmacodynamic functions: a multiparameter approach to the design of antibiotic treatment regimens. Antimicrob. Agents Chemother. 48, 3670–3676 (2004).

19. Abraham, M. J. et al. GROMACS: High performance molecular simulations through multi-level parallelism from laptops to supercomputers. SoftwareX 1, 19–25 (2015).

20. Best, R.B. et al. Optimization of the additive CHARMM all-atom protein force field targeting improved sampling of the backbone φ, ψ and side-chain χ1 and χ2 dihedral angles. J. Chem. Theory Comput. 8 (9), 3257–3273 (2012).

21. Huang, J., & MacKerell, A.D. CHARMM36 all-atom additive protein force field: validation based on comparison to NMR data. J. Comput. Chem. 34, 2135–2145 (2013).

22. Lee, J. et al. CHARMM-GUI input generator for NAMD, GROMACS, AMBER, OpenMM, and CHARMM/OpenMM simulations using the CHARMM36 additive force field. J. Chem. Theory Comput. 12 (1), 405–413 (2016).

23. Epand, R.M. & Epand, R.F. Bacterial membrane lipids in the action of antimicrobial agents. J. Pept. Sci. 17, 298–305 (2011).

24. Sohlenkamp, C. & Geiger, O. Bacterial membrane lipids: diversity in structures and pathways. FEMS Microbiology Reviews 40, 133–159 (2016).

25. Angelova, M. & Dimitrov, D.S. Liposome electroformation. Faraday Discuss. Chem. Soc. 81, 303–311 (1986).

26. Angelova, M. & Dimitrov, D.S. A mechanism of liposome electroformation. Trends in Colloid and Interface Science II, ed Degiorgio V (Springer, Berlin), 59–67 (1988).

27. Angelova, M. Giant vesicles. Perspectives in Supramolecular Chemistry, eds Luisi PL, Walde P (Wiley-Interscience, Chichester, UK), 1st Ed, 27–36 (2000).

28. Lindsey, H., Petersen, N.O. & Chan, S.I. Physicochemical characterization of 1,2-diphytanoyl-sn-glycero-3-phosphocholine in model membrane systems. Biochim. Biophys. Acta 555, 147–167 (1979).

29. Redwood, W.R., Pfeiffer, F.R., Weisbach, J.A. & Thompson, T.E. Physical properties of bilayer membranes formed from a synthetic saturated phospholipid in n-decane. Biochim. Biophys. Acta 233, 1–6 (1971).

30. Tosatto, L. et al. Alpha-synuclein pore forming activity upon membrane association. Biochim. Biophys. Acta 1818, 2876–2883 (2012).

31. Manzo, G., Hind, C.K., Ferguson, P.M., Amison, R.T, Hodgson-Casson, A.C., Ciazynska, K.A., Weller, B.J., Clarke, M., Lam C., Man, R.C.H., O’Shaughnessy, B.G., Clifford, M., Bui, T.T., Drake, A.F., Atkinson, R.A., Lam, J.K.W., Pitchford, S.C., Page, C.P., Phoenix, D.A., Lorenz, C.D., Sutton, J.M. & Mason, A.J. A Pleurocidin analogue with greater conformational flexibility, enhanced antimicrobial potency and in vivo therapeutic efficacy. Communications Biology 3, 697. (2020).

32. Torrico, M., Giménez, M.J., González, N., Alou, L., Sevillano, D., Cafini, F., Prieto, J., Cleeland, R & Aguilar, L. Bactericidal activity of daptomycin versus vancomycin in the presence of human albumin against vancomycin-susceptible but tolerant methicillin-resistant *Staphylococcus aureus* (MRSA) with daptomycin minimum inhibitory concentrations of 1-2 microg/mL. Int. J. Antimicrob. Agents 35, 131–137 (2010).

33. Zhanel, G.G., Voth, D., Nichol, K., Karlowsky, J.A., Noreddin, A.M. & Hoban, D.J. Pharmacodynamic activity of ceftobiprole compared with vancomycin versus methicillin-resistant *Staphylococcus aureus* (MRSA), vancomycin-intermediate *Staphylococcus aureus* (VISA) and vancomycin-resistant *Staphylococcus aureus* (VRSA) using an *in vitro* mode. J. Antimicrob. Chemother. 64, 364–369 (2009).

34. Manzo, G., Gianfanti, F., Hind, C.K., Alison, L., Clarke, M., Hohenbichler, J., Limantoro, I., Martin, B., Do Carmo Silva, P., Ferguson, P.M., Hodgson-Casson, A.C., Fleck, R.A., Phoenix, D.A. & Mason, A.J. Impacts of metabolism and organic acids on cell wall composition and *Pseudomonas aeruginosa* susceptibility to membrane active antimicrobials. ACS Infect. Diseases 7, 2310–2323 (2021).

35. Lipkin, R. & Lazaridis, T. Computational studies of peptide-induced membrane pore formation. Phil. Trans. R. Soc. B 372: 20160219 (2017).

36. Leontiadou, H., Mark, A. E. & Marrink, S. J. Antimicrobial peptides in action. J. Am. Chem. Soc. 128, 12156–12161 (2006).

37. Sengupta, D., Leontiadou, H., Mark, A. E. & Marrink, S. J. Toroidal pores formed by antimicrobial peptides show significant disorder. Biochim. Biophys. Acta 1778, 2308–2317 (2008).

38. Amos, S-B.T.A. et al. Antimicrobial peptide potency is facilitated by greater conformational flexibility when binding to Gram-negative bacterial inner membranes. Sci. Rep. 6, 37639–37651 (2016).

39. Mangoni, M.L., Di Grazia, A., Cappiello, F., Casciaro, B. & Luca, V. Naturally occurring peptides from *Rana temporaria*: Antimicrobial properties and more. Curr. Top. Med. Chem. 16, 54–64 (2016).

40. Rinaldi, A.C. et al. Temporin L: antimicrobial, haemolytic and cytotoxic activities, and effects on membrane permeabilization in lipid vesicles. Biochem. J. 368, 91–100 (2002).

41. Mangoni, M.L. et al. Effects of the antimicrobial peptide temporin L on cell morphology, membrane permeability and viability of *Escherichia coli*. Biochem. J. 380, 859–865 (2004).

42. Mangoni, M.L. et al. Structure-activity relationship, conformational and biological studies of temporin L analogues. J. Med. Chem. 54, 1298–1307 (2011).

43. Merlino, F. et al. Glycine-replaced derivatives of [Pro^3^,DLeu^9^]TL, a temporin L analogue: Evaluation of antimicrobial, cytotoxic and hemolytic activities. Eur. J. Med. Chem. 139, 750–761 (2017).

44. Saviello, M.R. et al. New insight into the mechanism of action of the temporin antimicrobial peptides. Biochemistry 49, 1477–1485 (2010).

45. Giacometti, A. et al. Interaction of antimicrobial peptide temporin L with lipopolysaccharide *in vitro* and in experimental rat models of septic shock caused by Gram-negative bacteria. Antimicrob. Agents Chemother. 50, 2478–2486 (2006).

46. Srivastava, S. & Ghosh J.K. Introduction of a lysine residue promotes aggregation of temporin L in lipopolysaccharides and augmentation of its antiendotoxin property. Antimicrob. Agents Chemother. 57, 2457–2466 (2013).

47. Srivastava, S., Kumar, A., Tripathi, A.K., Tandon, A. & Ghosh, J.K. Modulation of anti-endotoxin property of temporin L by minor amino acid substitution in identified phenylalanine zipper sequence. Biochem. J. 473, 4045–4062 (2016).

48. Wade, D., Silberring, J., Soliymani, R., Heikkinen, S., Kilpeläinen, I., Lankinen, H. & Kuusela P. Antibacterial activities of temporin A analogs. FEBS Lett. 479, 6–9 (2000).

49. Grassi, L., Maisetta, G., Maccari, G., Esin, S. & Batoni, G. (2017) Analogs of the frog-skin antimicrobial peptide Temporin 1Tb exhibit a wider spectrum of activity and a stronger antibiofilm potential as compared to the parental peptide. Front. Chem. 5: 24. doi: 10.3389/fchem.2017.00024 (2017).

50. Bhunia, A., Saravanan, R., Mohanram, H., Mangoni, M.L. & Bhattacharjya, S. NMR structures and interactions of temporin-1Tl and temporin-1Tb with lipopolysaccharide micelles: mechanistic insights into outer membrane permeabilization and synergistic activity. J. Biol. Chem. 286, 24394–24406 (2011).

51. Park, C.B., Kim, H.S. & Kim, S.C. Mechanism of action of the antimicrobial peptide buforin II: buforin II kills microorganisms by penetrating the cell membrane and inhibiting cellular functions. Biochem. Biophys. Res. Commun. 244, 253–247 (1998).

52. Park, C.B., Yi, K-S., Matsuzaki, K. Kim, M.S. & Kim, S.C. Structure–activity analysis of buforin II, a histone H2A-derived antimicrobial peptide: The proline hinge is responsible for the cell-penetrating ability of buforin II. Proc. Natl. Acad. Sci. USA 97, 8245–8250 (2000).

53. Lan, Y., Ye, Y., Kozlowska, J., Lam, J.K.W., Drake, A.F. & Mason, A.J. Structural contributions to the intracellular targeting strategies of antimicrobial peptides. Biochim. Biophys. Acta 1798, 1934–1943 (2010).

54. Yu, G., Baeder, D.Y., Regoes, R.R. & Rolff, J. Combination effects of antimicrobial peptides. Antimicrob. Agents Chemother. 60, 1717–1724 (2016).

55. Strandberg, E., Zerweck, J., Horn, D., Pritz, G., Berditsch, M., Bürck, J., Wadhwani, P. & Ulrich, A.S. Influence of hydrophobic residues on the activity of the antimicrobial peptide Magainin 2 and its synergy with PGLa. J. Pept. Sci. 21, 436–445 (2015).

56. Zerweck, J., Strandberg, E., Bürck, J., Reichert, J., Wadhwani, P., Kukharenko, O. & Ulrich, A.S. Homo- and heteromeric interaction strengths of the synergistic antimicrobial peptides PGLa and Magainin 2 in membranes. Eur. Biophys. J. 45, 535–547 (2016).

57. Zerweck, J., Strandberg, E., Kukharenko, O., Reichert, J., Bürck, J., Wadhwani, P., Ulrich, A.S. Molecular mechanism of synergy between the antimicrobial peptides PGLa and Magainin 2. Sci. Rep. 7, 13153 (2017).

58. Aisenbrey, C., Amaro, M., Pospíšil, P., Hof, M. & Bechinger, B. Highly synergistic antimicrobial activity of Magainin 2 and PGLa peptides is rooted in the formation of supramolecular complexes with lipids. Sci. Rep. 10, 11652 (2020).

59. Ma, W., Sun, S., Li, W., Zhang, Z., Lin, Z., Xia, Y., Yuan, B. & Yang, K. Individual roles of peptides PGLa and Magainin 2 in synergistic membrane poration. Langmuir 36, 7190–7199 (2020).

60. White, S.H. & Wimley, W.C. Hydrophobic interactions of peptides with membrane interfaces. Biochim. Biophys. Acta 1376; 339–352 (1998).

