## Supplementary Figures 1-7 for "Temporin B forms hetero-oligomers with Temporin L, modifies its membrane activity and increases the cooperativity of its antibacterial pharmacodynamic profile"

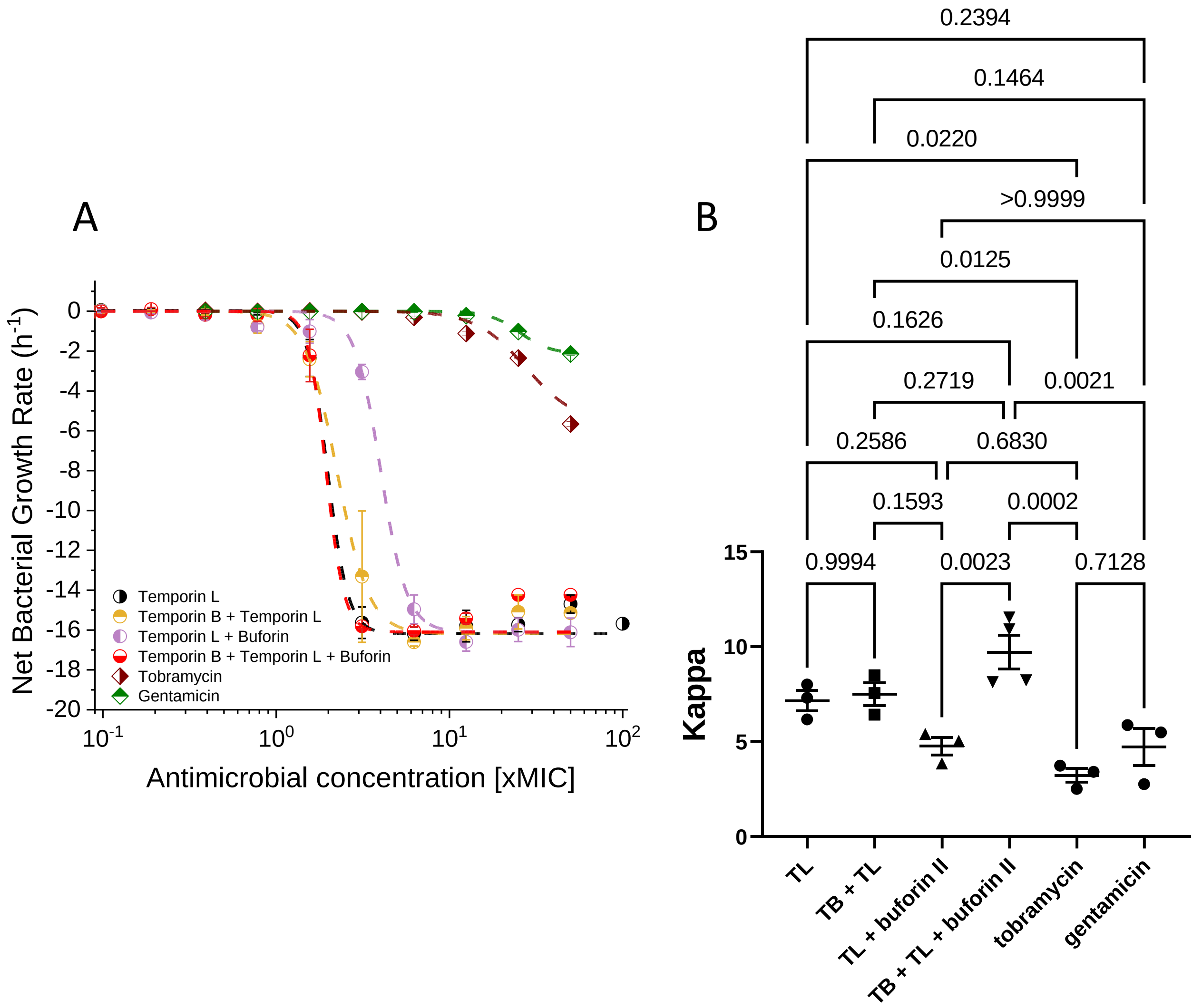


| Condition | Kappa | zMIC (xMIC) | $\psi_{max}$ ($h^{-1}$) | $\psi_{min}$ ($h^{-1}$) |
|---|---|---|---|---|
| **Temporin L** | 7.16 ± 0.53 | 0.75 ± 0.36 | 0.12 ± 0.06 | -16.3 ± 0.3 |
| **Temporin L / Temporin B** | 7.49 ± 0.60 | 0.85 ± 0.53 | 0.03 ± 0.03 | -16.6 ± 0.2 |
| **Temporin L / Buforin II** | 4.76 ± 0.47 | 0.60 ± 0.26 | 0.02 ± 0.02 | -16.7 ± 0.5 |
| **Temporin L/ Temporin B / Buforin II** | 9.71 ± 0.89 | 0.22 ± 0.09 | 0.00 ± 0.00 | -15.9 ± 0.2 |
| **Tobramycin** | **3.22 ± 0.37** | 1.05 ± 0.36 | 0.00 ± 0.00 | **-5.66 ± 0.14** |
| **Gentamicin** | 4.70 ± 0.98 | 2.08 ± 0.65 | 0.00 ± 0.00 | **-2.14 ± 0.08** |

**Figure S1. Pharmacodynamic response of EMRSA-15 to antibiotic challenge in LB.** EMRSA-15 was challenged with increasing concentrations of temporin L (TL), a 16:1 mol:mol ratio combination of temporin B and temporin L (TB:TL) or a 16:1:16 mol:mol:mol combination of temporin B, temporin L and buforin II. Curves shown are fits of averages of three independent repeated experiments (A). The cooperativity (Kappa), pharmacodynamic MIC (zMIC) and maximum ($\psi_{max}$) and minimum ($\psi_{min}$) growth rates are provided in the table and were obtained by averaging fits of three or more independently repeated experiments. One-way ANOVA with Tukey post-hoc test multiple comparisons for Kappa, highlight the effect of adding all three peptides in combination (B).

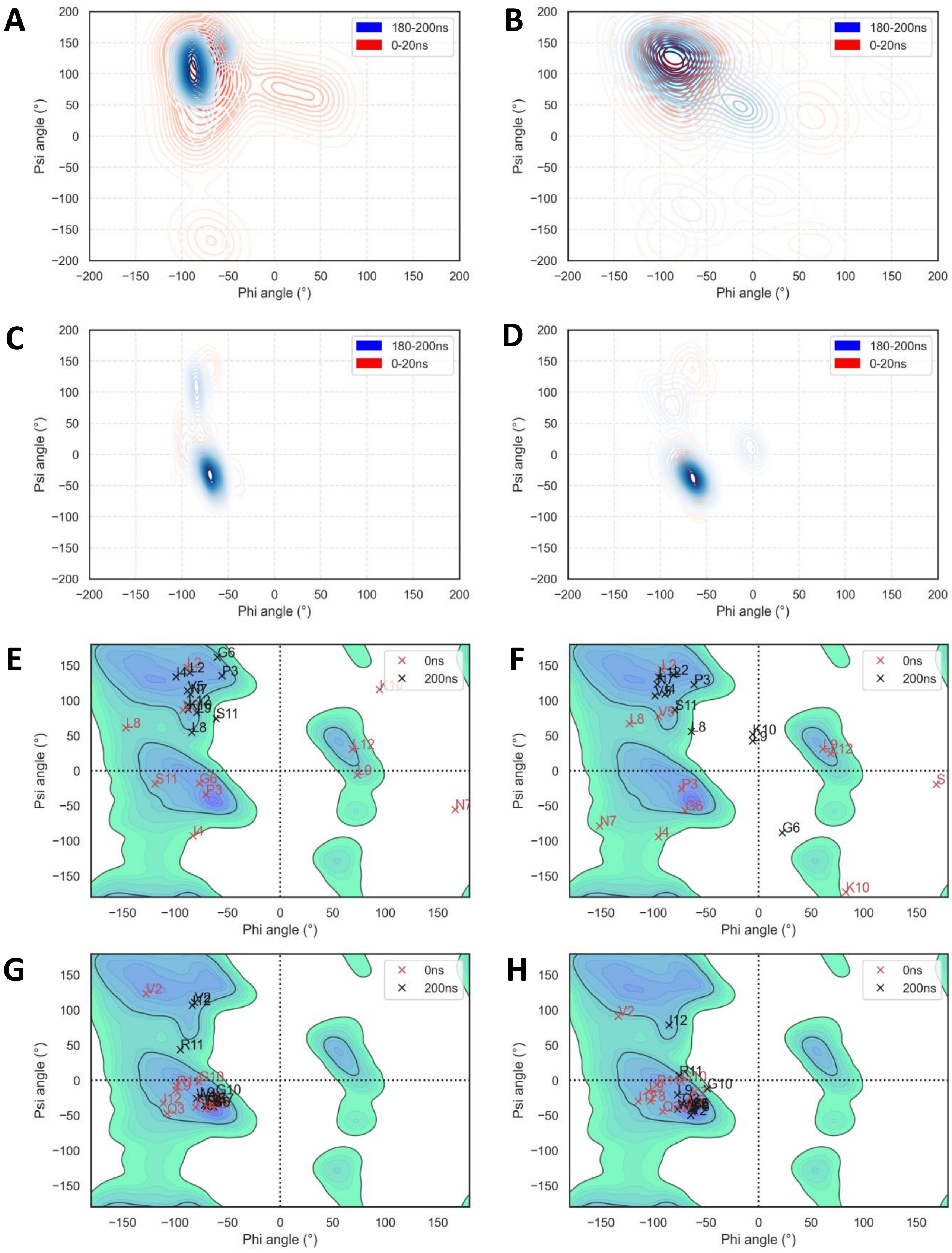


**Figure S2. Modest changes in peptide conformation when temporin B is added to temporin L in MD simulations of POPG bilayer challenge.** Ramachandran plots for eight temporin B peptides (A, E), temporin B in 4:4 combination with temporin L (B, F), eight temporin L peptide (C, G) and temporin L in 4:4 combination with temporin B (D, H). Contour plots (A-D) show dihedral angles averaged over time and over the peptide primary sequence while snapshots (E-H) show the dihedral angles for individual residues at the beginning and end of the simulation. Data is representative of two independently repeated simulations in each case.

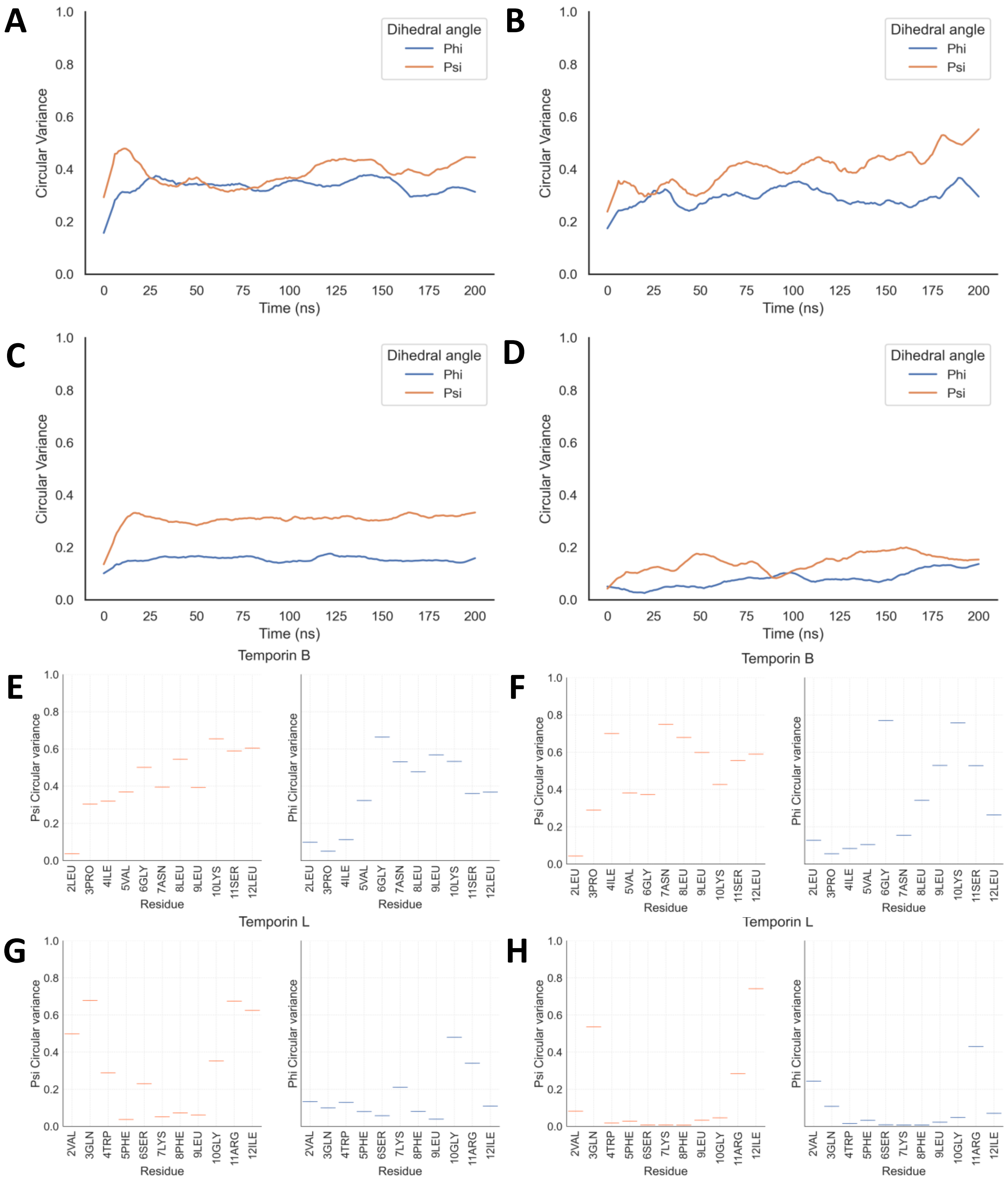


**Figure S3. Reduced peptide conformational flexibility for temporin L when temporin B is added in MD simulations of POPG bilayer challenge.** Dihedral angle circular variance is shown as an average across all residues (A-D) or as an average over the simulation duration as a function of residue position (E-H). These are provided for eight temporin B peptides (A, E), temporin B in 4:4 combination with temporin L (B, F), eight temporin L peptides (C, G) and temporin L in 4:4 combination with temporin B (D, H). Data is representative of two independently repeated simulations for each condition.

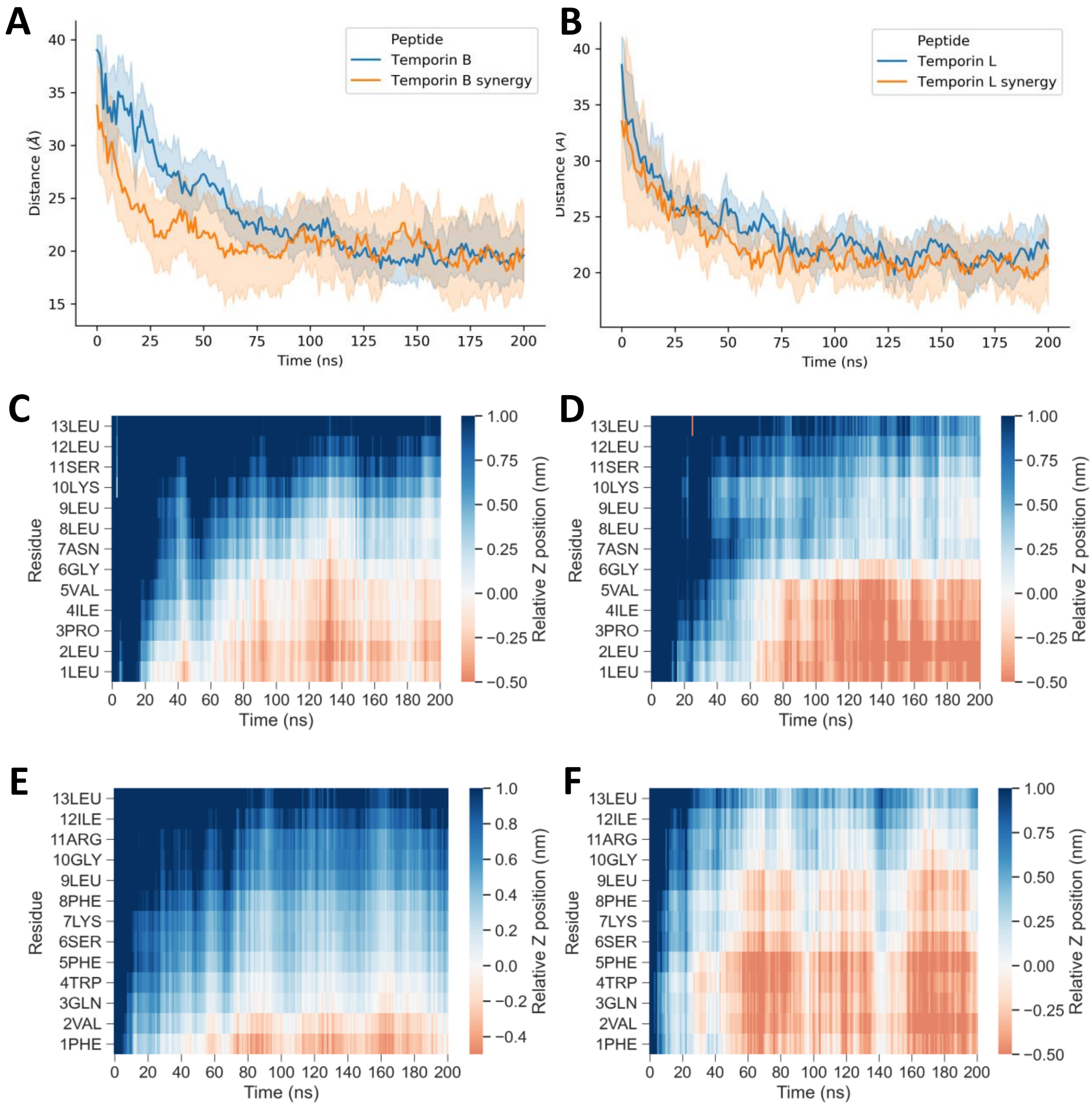


**Figure S4. MD simulations of temporin L and temporin B challenge of POPG bilayer.** Peptide insertion centre of mass over time for temporin B (A) or temporin L (B) comparing either eight temporin B or eight temporin L peptides with a 4:4 combination of each, focussing on the indicated peptide. Data is an average of two independently repeated simulations for each condition. Peptide penetration over time by primary sequence for eight temporin B peptides (C), temporin B in a 4:4 combination with temporin L (D), eight temporin L peptides (E) and temporin L in a 4:4 combination with temporin B (F). The Z-position is calculated for each residue, averaged over all eight or all four peptides, relative to the phosphate group plane in the upper bilayer leaflet. Data is representative of two independently repeated simulations for each condition.

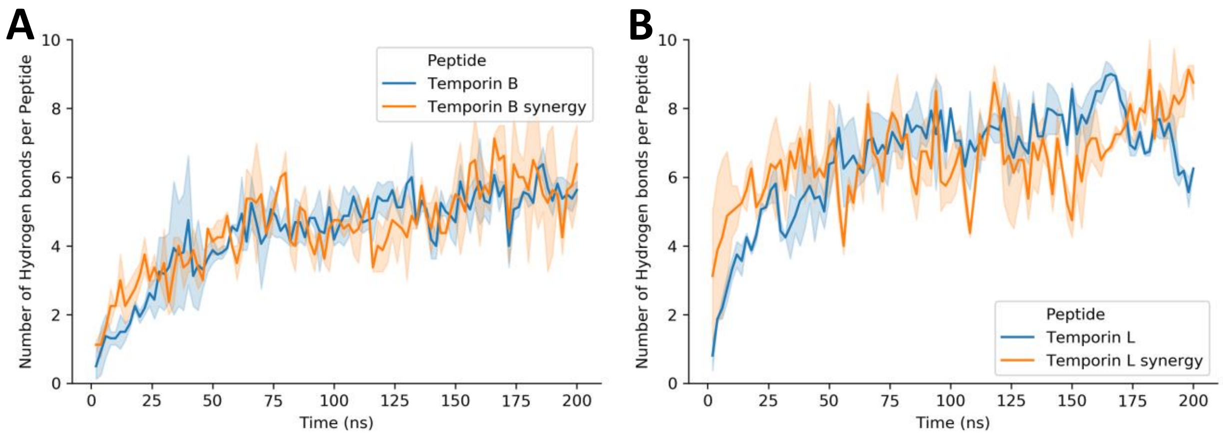


**Figure S5. MD simulations of temporin L and temporin B challenge of POPG bilayer.** Total number of peptide-lipid hydrogen bonds, per peptide, over time for eight temporin B (A), or temporin L (B), peptides binding to a 512-lipid bi-layer compared with the corresponding data for the peptides in a 4:4 combination. Averages ± standard error of two replicate simulations are shown.

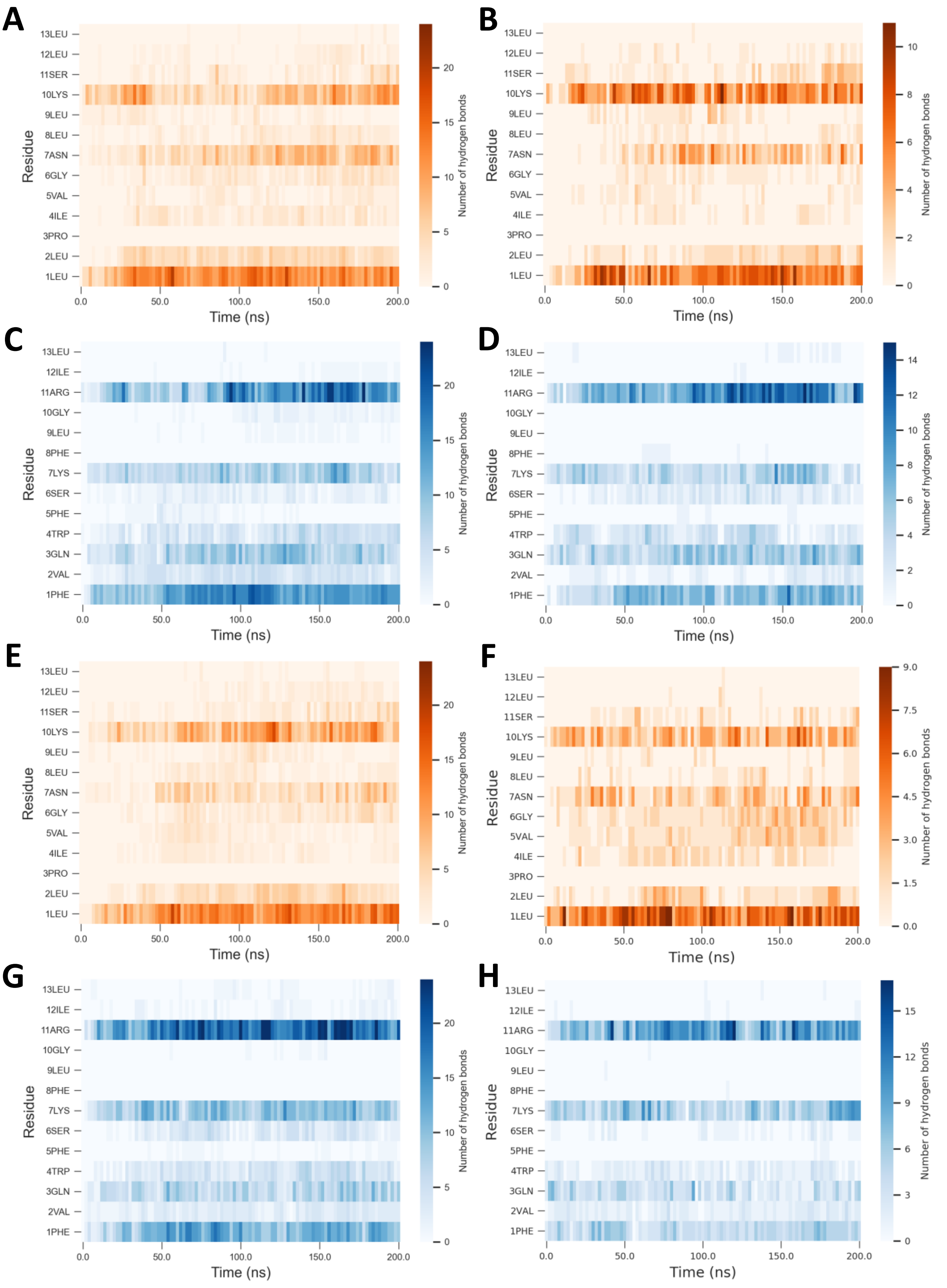


**Figure S6. Peptide-lipid hydrogen bonding pattern is unaltered by combining temporin L and temporin B.** Total, peptide-lipid hydrogen bonds by residue for duplicate simulations of: eight temporin B peptides (A/E); temporin B in a 4:4 combination with temporin L (B/F); eight temporin L peptide (C/G); and temporin L in a 4:4 combination with temporin B (D/H).

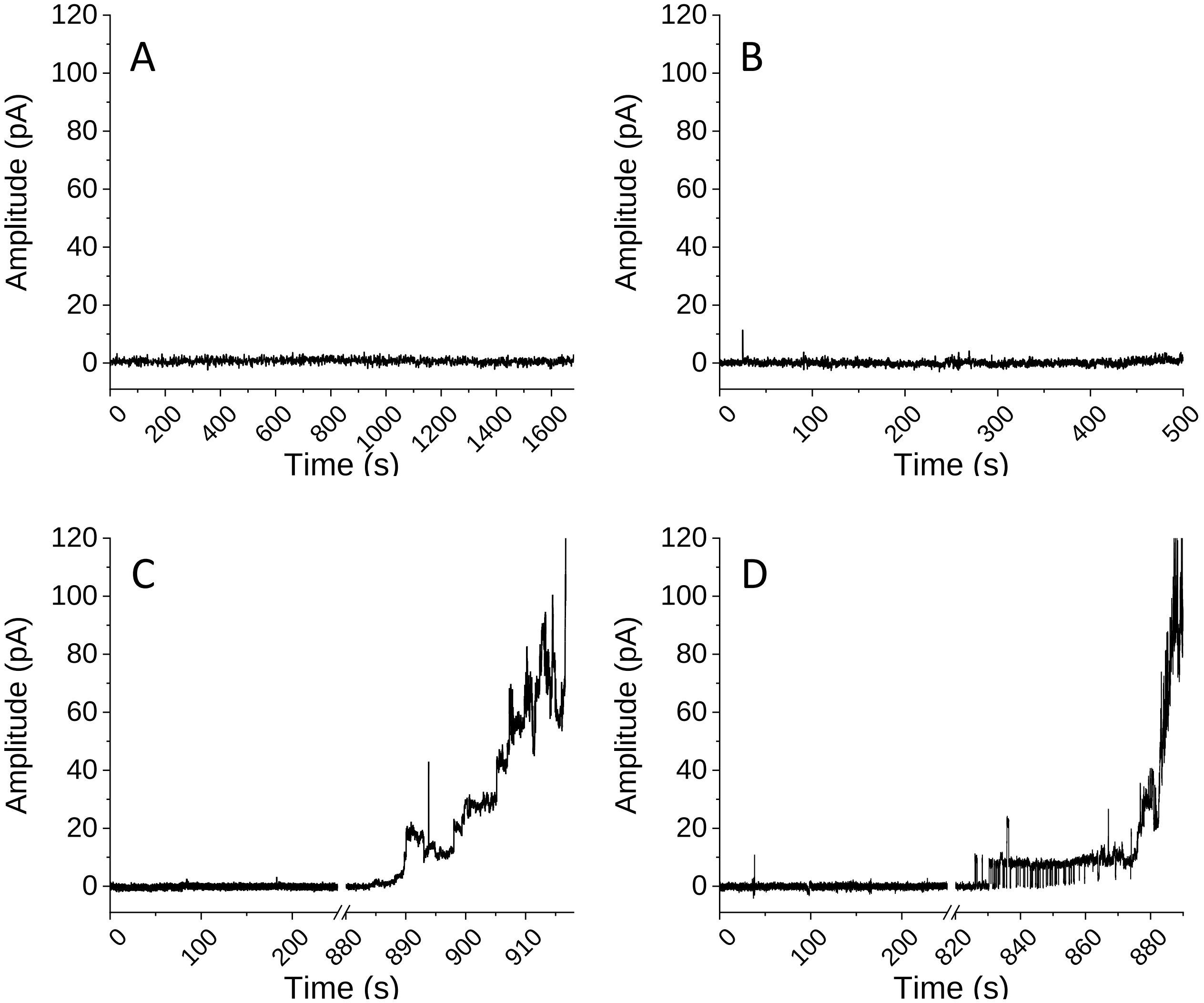


**Figure S7. Patch-clamp threshold concentration determination.** Representative traces show the absence of conductance below the threshold concentrations for temporin L (A) and the combination of temporin L and temporin B (B). No conductance is observed when membranes are challenged with 5 µM temporin L, ½ the threshold concentration (A). At its threshold concentration of 10 µM conductance is observed 1040 ± 86 seconds after administration of the peptide. No conductance is observed when membranes are challenged with 1.46 µM temporin B and 0.42 µM temporin L, ½ the threshold concentration (B). At the threshold concentration for the combination, conductance is observed at 432 ± 61 seconds after administration of the peptide. At double (5.84 µM temporin B and 1.68 µM temporin; C) or three times (8.75 µM temporin B and 2.5 µM temporin L; D) the threshold concentration for the combination higher amplitude conductance is obtained but this takes longer to begin.
